# Multiple forms of neural processing when repeating voluntary thumb flexions

**DOI:** 10.1101/2023.02.19.529148

**Authors:** Ruchella Kock, Arko Ghosh

**Author notes:** Corresponding address Arko Ghosh, Leiden University, Wassenaarseweg 52, Leiden, 2333 AK. **Author contributions** A.G. conceived the study. A.G. and R.K. designed the study. R.K. analyzed the data aided by A.G. A.G. drafted the report aided by R.K. Both the authors helped edit the manuscript. **Funding** This study was funded by a research grant from Velux Stiftung (no. 1283, A.G. is principal investigator). **Competing interests** None to declare.

## Abstract

There is considerable trial-to-trial variability in single cortical neurons when performing the same action repeatedly. One possibility is that neural populations are more stable in representing actions; alternatively, they too may be distinctly engaged from trial-to-trial. To address the nature of variability in large neural populations, we captured the EEG signals time-locked to repeated voluntary thumb flexion movements (∼500 repetitions, 23 subjects). By using non-negative matrix factorization on the low-frequency sensorimotor cortical signals, we quantified the trial-to-trial motor-related potentials (MRPs) in terms of prototypical signals (meta-MRPs) and their corresponding prominence at each trial (meta-trials). Clustering the meta-MRPs across the sampled population revealed 5 distinct signal patterns. There were brain-wide correlates of these meta-MRP clusters. Cortical hemispheres were distinctly recruited from trial-to-trial as certain clusters were accompanied by bilateral motor negativity while others were characterized by ipsilateral motor negativity. The sensory feedback too was distinctly processed from trial-to-trial as the central post-movement positivity was present only for certain clusters. A poorly understood pre-motor positivity accompanied all clusters albeit varying in their timing from trial-to-trial. These patterns – including the time-varying positivity preceding the movement – were rendered invisible by the traditional averaging of the signals. We suggest that the variability in EEG signals is not just noise but a consequence of distinct activation patterns deployed by the cortex. We support the idea that the cortex flexibly switches between distinct forms of neural processing to achieve the same behavioral goals.

## Introduction

Variability is ubiquitous in neural measures. This is vividly demonstrated in single-neuron recordings of the motor cortex that show trial-to-trial variability even in the absence of any behavioral variability (Lee et al., 1998); for review see (Renart and Machens, 2014). According to one idea, this variability is accompanied by redundant information processing – especially in the neighboring neurons – and the population-wide signals offer a more stable representation of movements (Carmena et al., 2005). Trial-to-trial variability is also observed in EEG – which captures the combined activity of large populations of neurons – may it be when actively performing actions or passively processing identical sensory inputs. According to a common assumption, much of this variability stems from measurement noise, and therefore, the underlying signals are inferred by simply averaging across the trials (Luck, 2014). An emerging body of work suggests that variability is not simply measurement noise as the amount of variability is different in people with clinical disorders such as autism in contrast to healthy controls (Dinstein et al., 2015). Still, the nature of the neural trial-to-trial variability captured in EEG is unclear. Are the same populations activated but at distinct magnitudes or are distinct populations activated from one trial-to-the next or both?

Inter-individual differences in EEG signals even performing rather simple voluntary actions suggests there can be multiple patterns of population activity for the same action, albeit it does not clarify the nature of trial-to-trial variations. First, the hallmark of movement preparation i.e., the slow pre-motor build-up of neural activity (negativity) prior to the movement onset is inexplicably absent in certain individuals (Schurger et al., 2021). Second, a positivity preceding the movement – with unclear function – is only visible in a fraction of the subjects (Ball et al., 1999). Finally, there are substantial inter-individual differences in the performance of brain computer interface algorithms used to infer simple finger movements from EEG signals (Wang et al., 2020).

Conventionally, event-related potentials (ERPs) are derived from a second order tensor of trials × amplitude at a given time, and the averaged signals are well understood in terms of their cognitive correlates. For instance, the ERPs time locked to the repeated movements (motor related potentials, MRPs) can be used to monitor the different stages of motor preparation and post-movement processing based on the signal timing and shape (Tarkka and Hallett, 1990). Still, this comes at the cost of the loss of information at the level of each trial. Dimensionality reduction of the tensor – without averaging – may help discover the hidden patterns that may well vary from trial-to-trial. The widely used principle component analysis, can reduce the tensor but it forces the components to be orthogonal to each other – making them difficult to interpret. In contrast, non-negative matrix factorization (NNMF) learns the parts-based representation and provides a more interpretable low-rank approximation of a tensor. For instance, an image containing a face is decomposed into features containing eyebrows, nose, etc. using NNMF but not when using PCA (Lee and Seung, 1999). In biology, NNMF method has been used to condense large gene expression data to a few *meta-genes* to help identify disease mechanisms (Brunet et al., 2004). Furthermore, NNMF is commonly used to identify muscle groups based on electromyography (EMG) recordings (Rabbi et al., 2020). In brain computer interface research, NNMF has been successfully applied to select the relevant electrode or to generate features for classifiers, but these efforts are not designed to unravel the nature of trial-to-trial variability when repeating actions (Lu and Yin, 2015; Mørup et al., 2007).

Here we applied NNMF to the low frequency (δ) sensorimotor EEG signal second order tensor (trials × amplitude at a given time) surrounding voluntary movements to determine a set of *meta-MRPs* and the corresponding *meta-trials*. Low-frequency EEG signals contain information on the motor-relevant object parameters. We limited the factorization to signals starting 500 ms before the completion of the thumb flexion and 100 ms after the flexion to focus on the variability in the motor-related signals. The meta-MRPs were the time-variant prototypical MRP signal features scattered across the trials and the meta-trials captured the prominence of the corresponding feature at a given trial. Subsequently, we clustered the meta-MRPs across the sampled population to identify the distinct signals captured from one-trial to the next. We leverage the corresponding meta-trials to discover the brain wide correlates that spanned EEG electrodes and time periods beyond motor processing.

## Results

### Conventional analysis of motor related potentials

Participants repeatedly performed voluntary thumb flexions to touch a rectangular box. The participants were instructed to touch the box at will, guided by a ∼ 5 s dial (**Figure 1**). By using a movement sensor attached on the thumb we determined the movement duration - sensor deflection to box contact – to be 164 ms (population median). We time-locked the EEG signals to the contact onset on the box; which we found to less equivocal than the movement onset due to the decisive signal on the force sensor (Supplementary Figure 1). The averaging of the EEG signals surrounding the movements revealed an early (at ∼-1s) component characterized by negativity over the central electrodes and positivity over the sensorimotor electrodes (Supplementary figure 2, Supplementary Movie 1). By - ∼500 ms the negativity was present bilaterally occupying the central and sensorimotor electrodes. The bilateral negativity persisted till about ∼-100 ms after which the signal became more lateralized to occupy the contralateral sensorimotor electrodes (to movement, i.e., left hemisphere). The contralateral negativity persisted till ∼150 ms after the contact onset. Subsequently, we observed a central positivity that lasted till about ∼300 ms after which it was replaced with a component that spanned the sensorimotor (negative), central and occipital electrodes (positive). This late component persisted till ∼650 ms after which weak signals were observed over the occipital electrodes. In sum, as anticipated based on prior work, the thumb flexions were accompanied by an early bilateral negativity, followed by a dominatingly contralateral negativity during the movement and a post-movement positivity.

**Figure 1.**
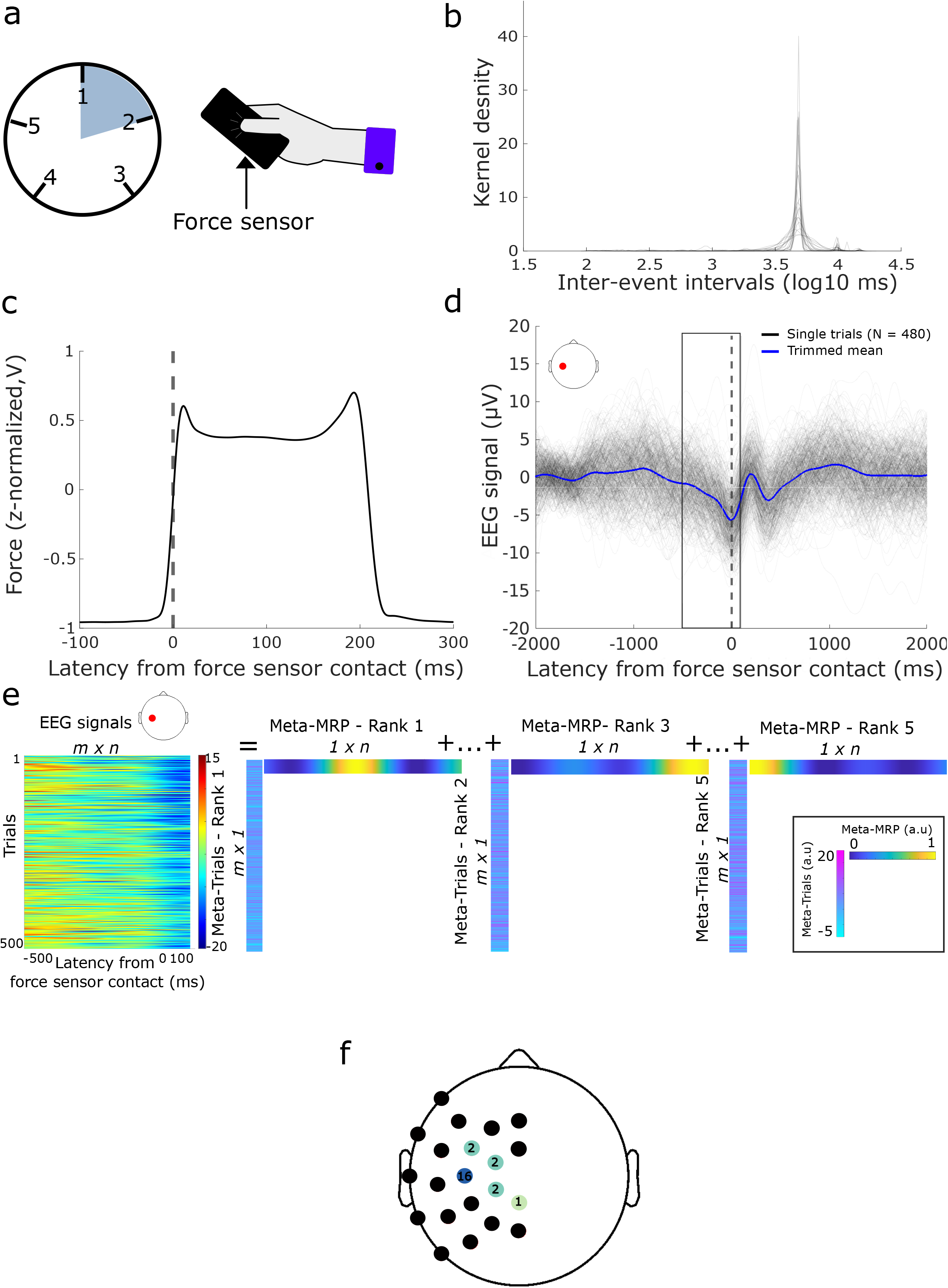
Methods towards identifying multiple neural strategies. **(a)** Illustration of experimental setup. Participants performed thumb flexions on a force sensor within a 5-second window shown as an analog clock. **(b)** Probability density of inter-event-intervals across the sampled population, showing that most touches occurred within around 3 seconds (log_10_ normalized for visualization). **(c)** Grand average force sensor contact signal time-locked to the end of the thumb flexions with a duration of 200 (+/- 20) ms (z-normalized for visualization). **(d)** A negative deviation in the event-related potential, over the left sensorimotor cortex, indicates preparation of the thumb flexions for one participant (blue line, trimmed means 20%). This starts at about ∼500 ms preceding the event (black outlined box), and is followed by a positive deflection indicating processing of the movement. Single trials (black lines) demonstrate trial-by-trial variability. **(e)** Applying non-negative matrix factorization to a contralateral sensorimotor electrode of one participant. Two-dimensional time-locked EEG data (−500 to 100 ms) is decomposed in the optimal number of factorizations (for this subject, 5 ranks, only Rank 1,3 and 5 are shown). Each rank consists of two vectors representing prototypical motor-related potentials (meta-MRPs) and similarity of each trial to the meta-MRPs (meta-Trials). The meta-MRPs demonstrate clear differences in the preparation of the thumb flexion. For instance, rank 3 shows a strong positive deflection near the touch, while rank 5 is categorized by a negative deflection near the touch. **(f)** Selection of sensorimotor electrodes based on the Surface Laplacian technique.

### Prototypical motor related potentials based on NNMF (meta-ERPs)

NNMF was used to reduce the EEG signals captured from a select electrode over the sensorimotor cortex and time-locked to the contact onset. The tensor (trials × amplitude at a given time between - 500 ms to 100 ms) was reduced to a few of meta-MRPs (i.e., the prototypical signal features) and the corresponding meta-trials (the prominence of the reduced features at a given trial, **Figure 1**). Meta-MRPs from the sampled population were grouped into 5 clusters according to their shape (**Figure 2**). The most common meta-MRP cluster (contributed by all of the subjects) was characterized by a signal peaking at the time of the contact. The next common cluster (contributed by 16 of the 23 subjects) was characterized by an early signal peaking at ∼-400 ms. The third cluster (contributed by 12 of the 23 subjects) showed a signal peaking at ∼-200 ms prior to the contact. The fourth cluster (contributed by 11 of the 23 subjects) was characterized by a slow negative slope spanning the - 500 ms to + 100 ms factorization window. The final cluster showed a signal peaking at ∼-100 ms (contributed by 10 of the 23 subjects). In sum, NNMF extracted distinct prototypical signals from single trial EEG and these patterns were common in the sampled population.

**Figure 2.**
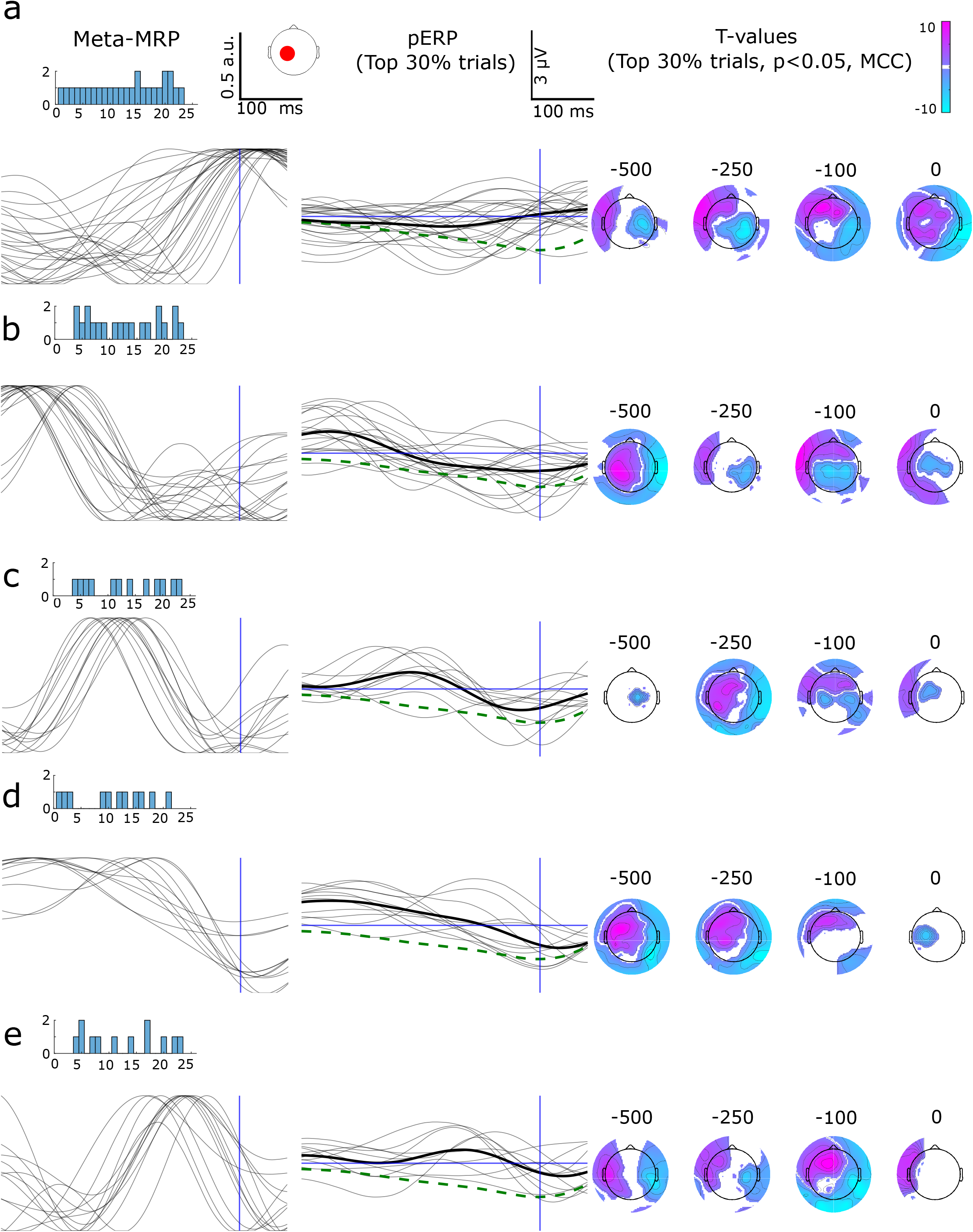
Non-negative matrix factorization applied to EEG signals resulted in 5 consistent clusters of neural activity across the population. Meta-MRPs reveal consistent patterns across the population (left column). The participants in each cluster are shown with a bar plot. Subsequently, the time-locked EEG trials (−500 to 100 ms from the end of the thumb flexion) were sorted, with the meta-trials, by similarity to the meta-MRPs. The top 30 percent of the sorted trials were used to generate the event-related potentials (pERPs) of the electrode located over the sensorimotor cortex (middle column, trimmed mean 20%). Through the clustered population event-related potentials (thick black line) it is clear that the selected trials follow a similar shape as the meta-MRPs, and that they differ from the grand average (dashed green lines). One participant’s pERPs was removed from the plots for visibility. Scalp topologies of T-values (top 30 percent of the selected trials) display statistically significant activity preceding the touch (right column). T-values are based on one-sample *t*-tests and multiple comparison corrections across time and electrodes (MCC, p < 0.05). Some clusters are dominated by bilateral motor negativity while others contralateral motor negativity. Each row represents one cluster. For the full statistical outcomes of these event-related potentials, see Supplementary Movie 2 to 6. For paired statistics of clusters, a and b, see Supplementary Figure 5 and Supplementary Movie 7.

### Variance of the meta-MRPs captured using meta-trials

The prominence of the meta-MRPs was captured in the corresponding meta-trials. The meta-trials varied substantially from one trial to the next. First, we addressed if the meta-trials systematically varied from one trial to the next. We reasoned that if the slow states (> 5 s) shaped the signals then the subsequent meta-trials would be related to each other. However, we observed no such relationship (Supplementary Figure 3). Second, we addressed whether the meta-trials were correlated to the trial-to-trial behavioral fluctuations. Towards this, we estimated the regression slopes for meta-trials vs. behavioral attribute pairs. The attributes included the force generated, gap from the previous behavioral event, the duration of the contact and the time from the movement onset. There were no consistent slopes for any of the meta-MRP clusters (the distribution of the coefficients is reported in Supplementary Figure 4).

### Brain-wide correlates of the meta-MRPs

Distinct set of ERPs (henceforth referred to as partial ERP, *pERP*) – spanning all the electrodes – were derived based on the top 30^th^ percentile of the meta-trials. The pERPs were then pooled together across the population according to the corresponding meta-MRP clusters followed by one sample *t*-tests (See Supplementary Movies 2 to 6, **Figure 2**). For all the five meta-MRP clusters the pERPs were characterized by a positive signal over the contralateral (to movement, i.e., the left hemisphere) sensorimotor cortex starting at about ∼1s prior to the contact event, albeit the positivity was stronger for some clusters than others (see paired statistics comparing the top two clusters in Supplementary Figure 5, and Supplementary Movie 7). The pERP based on the first cluster showed a unique pre-motor ipsilateral (to movement, i.e., right hemisphere) negativity between ∼-500 ms to ∼-250 ms. The second and third cluster showed an initially bilateral negativity followed by a contralateral negativity at contact. In the fourth cluster the sensorimotor negativity was mostly contralateral and appeared close to the contact time. The final cluster was dominated by ipsilateral negativity between ∼-350 to ∼-200 ms. These findings indicate that distinct neural populations were engaged for the neural control of the movement from trial-to-trial.

Post-movement, the distinct patterns of pERPs were observed for the meta-MRP clusters. The first cluster showed a prominent central positivity peaking at ∼150 ms (Fig.3, Supplementary Movie 2). In the second cluster the positivity was less pronounced (for paired analysis see Supplementary Figure 5, and Supplementary Movie 7). The third and the fourth clusters did not show any significant positivity, but the fourth cluster did show a late contralateral positivity. The fifth cluster showed a significant central positivity followed by a more diffuse component spanning the occipital electrodes as well.

**Figure 3.**
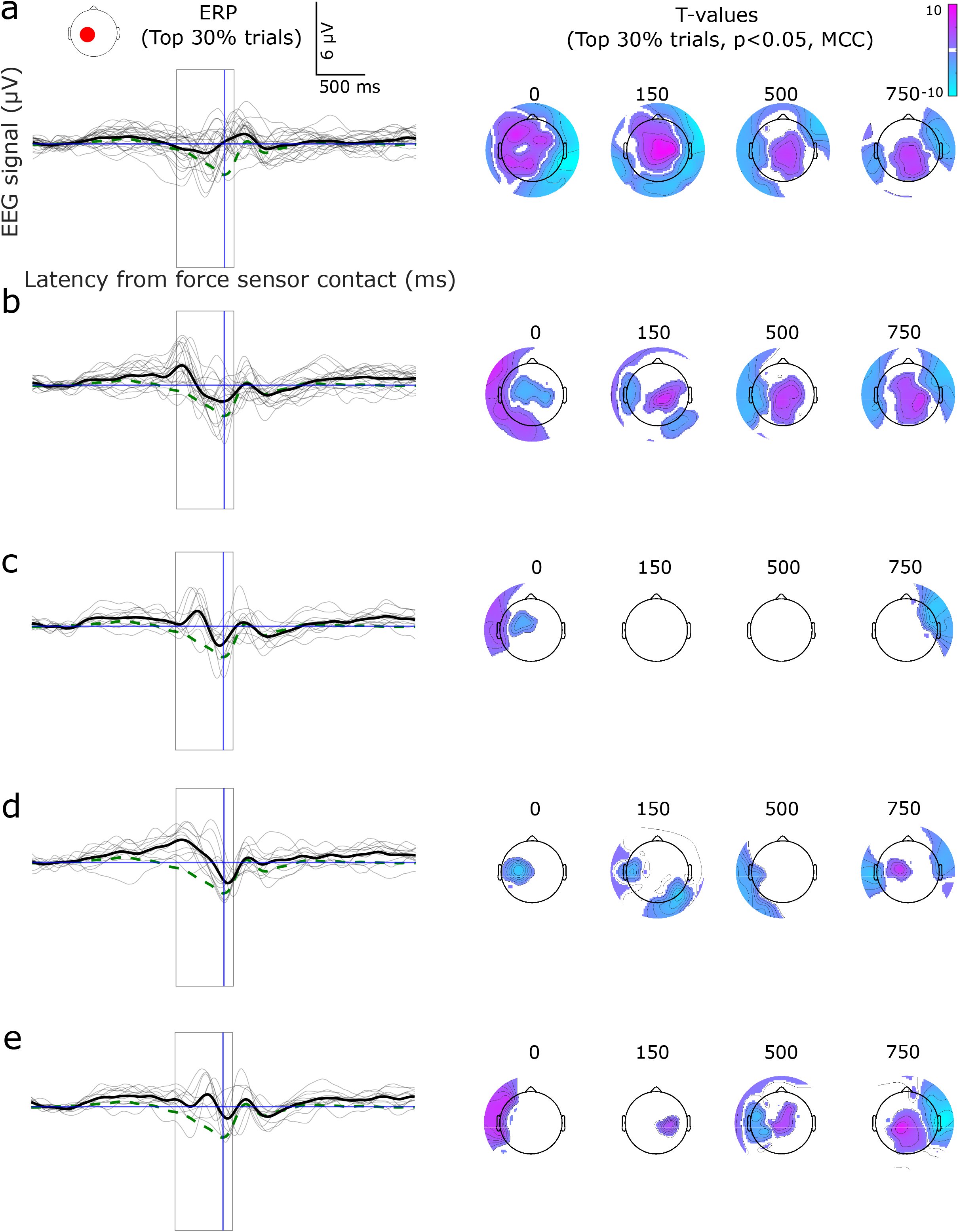
The clustered neural patterns are categorized by distinct neural processing even following the event. Event-related potentials for the sorted trials showing the full time period of the thumb flexion (left column, trimmed mean 20%). The grand average is shown through dashed green lines and clustered population average with black thick lines. The preparation time over which the non-negative matrix factorization was performed is shown in a black outlined box. Scalp topologies of T-values (top 30 percent of the selected trials) display statistically significant activity following the touch (right column, MCC, p < 0.05). Only certain clusters display a post movement positivity.

## Discussion

We captured distinct neural activation patterns embedded in the trial-to-trial low-frequency (δ band) EEG signal variations surrounding voluntary thumb flexions. These patterns – spanning various sensorimotor processing stages – were observed across the sampled population, albeit some of the patterns were more consistently observed than the others. The patterns differed considerably from each other in terms of the hemispheres engaged, the timing of the signals and the amplitude of the signals. The patterns were not simply related to the trial-to-trial behavioral differences.

Our findings help clarify the nature of neural variability surrounding voluntary movements. In theory, the variations could occur in the form of the same neural populations engaged but to generate different amounts of activity. For instance, attending to a stimuli can enhance the firing rate according to multi-unit recordings and increases the population level activity captured on EEG (Arazi et al., 2019; Noudoost et al., 2010). Our results suggest an alternative form of variability where entirely distinct populations can be engaged from trial-to-trial. According to our NNMF analysis certain trials primarily engaged the ipsilateral cortex for motor preparation and execution, other trials engaged the cortex bilaterally or contralaterally as indicated by the negativity in pERPs (which were based on the sorted meta-trials).

The inconsistent engagement of the ipsilateral cortex also challenges the traditional account of motor preparation based on the signal averages, according to which motor preparation occurs bilaterally followed by sustained engagement of the contralateral cortex for motor execution (also reported here using the averaged signals)(Colebatch, 2007; Deecke et al., 1976; Qing Cui and Deecke, 1999). The putative contribution of the ipsilateral cortex is distinct from the contralateral cortex – for instance, by being more engaged in complex tasks (Chen et al., 1997; Qing Cui and Deecke, 1999). While this may be the case, it does not help explain as to why certain trials show dominant ipsilateral activity (negativity) in our task. There may be redundant representations for movement distributed in the ipsilateral and contralateral cortex, and the representation used for motor control varies from trial-to-trial (Cauraugh and Summers, 2005; So et al., 2012).

The motor related positivity in the contralateral hemisphere – a less understood and inconsistently reported signal than the negativity associated with motor control (Shibasaki and Hallett, 2006) – was distinctly timed in the meta-ERP clusters and the corresponding pERPs. According to one idea this positivity indicates a *go* signal for motor execution (Bortoletto et al., 2006; Deecke et al., 1976). As the signals were time-locked to flexion endpoint it is possible that these differences stemmed from the trial-to-trial differences in the onset of the movement (in terms of the underlying muscular activation). However, we did not find evidence supporting this idea as the meta-trials were not related to the movement time (i.e., time gap between movement onset according to the movement sensor and the end of the thumb flexion). Another possibility is that the positivity indicates an inhibitory signal (Misirlisoy and Haggard, 2014). Following this interpretation, we speculate that the inhibitory processes may influence motor processing at different moments from trial-to-trial shaped by the overall network dynamics. For instance, at certain trials motor inhibition may have been evoked pre-emptively whereas in other trials it may have been to prevent ongoing muscular activity. The time variant nature of the positivity helps explain why they have been only inconsistently observed in previous literature (Karp et al., 1996; Tarkka and Hallett, 1990).

Although the NNMF focused on the signals surrounding the movement (−500 ms to 100 ms from the end of the thumb flexion), the pERPs revealed different patterns of post-movement processing from trial-to-trial accompanying the meta-MRPs. While certain pERPs showed strong post-movement positivity (150 to 750 ms) over the central and occipital electrodes, other pERPs did not show such significant statistical clusters. The post-movement positivity – also visible in averaged signal – putatively reflects the processing of sensory feedback from the movement (termed as reafferent potentials in (Kornhuber and Deecke, 2016)). The variations captured here may be related to the confidence of the feedforward estimation of the planned movement (Tan et al., 2016). Taken together with our findings from the motor period, perhaps certain movement representations result in distinct quality of feedforward estimations.

While there is support for the idea that single neurons contain redundant representations for movements. Our findings suggest that there may be redundant representations of movements in large neural populations as well. Moreover, the variations in single neurons do not simply relate to behavioral parameters. Here too we did not find behavioral correlates of the neural activation patterns. Our approach adds to single trial EEG analysis by enabling the exploration of inconsistent neural processing. The patterns discovered here encourage us to more broadly apply interpretable data-driven reduction of complex neural signals to unravel the key principles underlying brain functions.

## Methods

### Participants

Twenty-five right-handed participants were recruited at the Leiden University campus through flyers and advertisement. This recruitment was part of a larger study on smartphone behavior (Kock et al., 2022). The experiment was approved by the Psychology Ethics Committee and all participants provided written informed consent.

### Movements collected on a smartphone-shaped sensor

A hand-held, unresponsive device was created by placing a force resistive sensor (Interlink Electronics, Camarillo, USA) on a rectangular flat metal panel (further referred as force sensor) Participants performed voluntary thumb flexions on the device within a 5-second period, indicated by an analog clock. Only touches within the 2 mm target perimeter were recorded, and the sensor did not elicit any feedback. The force sensor output was recorded using a USB 6008 DAQ (National Instruments, Austin, USA, Figure 1a).

The force sensor output exhibited measurement noise, oscillating between -1 and -0.8 V, when no force was applied. To remove this noise during contact, we conducted Fast Fourier Transform on the sequences where no force was exerted. Predominant noise frequencies were below 0.05 Hz, and were removed using a lowpass filter.

The end of the thumb flexion was marked by a sudden rise in signal, followed by a slight release of force and a rapid drop to baseline (**Figure 1**). The participants periodically released the sensor for approximately 5 ms during certain touches. We merged the signals before and after the release (with an inter-event interval less than 5 ms), since they belonged to the same action. Furthermore, touches with inter-event-intervals shorter than 100 ms were excluded.

### EEG data collection and preprocessing

EEG data was collected while participants performed thumb flexions on the force sensor box. The measurement was performed using a 64-channel EEG cap equipped with equidistant electrodes (Easycap GmbH, Wörthsee, Germany) and ABRALYT HiCl electrode gel. The 64-channel DC amplifier BrainAmp (Brain Products GmbH, Gilching) was utilized for data acquisition, which was then digitized at a rate of 1 kHz. During the measurement, the participants were seated in a Faraday cage.

Processing of the EEG data included interpolating electrodes with impedances exceeding 10 kΩ and blink artifacts removal through Independent Component Analysis. For event-related analysis, the data was filtered between 0.5 and 3 Hz and segmented into -2 to 2 s epochs before and after the contact with the force sensor. The epoched data went through baseline correction, by subtracting the mean amplitude (over the interval of -2 s to -1.5 s) from the data. Trials with amplitudes exceeding ± 80 µV were rejected as artifacts. Two participants were excluded from the analysis due to insufficient trials after pre-processing (smaller than 200, Supplementary Figure 5). Additionally, the two ocular electrodes placed beneath the eyes were also removed from all further analysis.

### Electrodes selection

As the focus was on the thumb flexions, electrodes located near the sensorimotor cortex were selected. Given that participants were moving their right thumb, the selection space was constrained to 21 electrodes in the left parietal lobe. Next, the Surface Laplacian transformation was performed over the event-related potentials to enhance spatial resolution (Babiloni et al., 1995). For each electrode, the median over the event-related potential was calculated over a –500 to 100 ms window from the end of the thumb flexion. The electrode with the smallest median was selected for further analysis, as it reflected the negative signal associated with motor preparation (**Figure 1**).

### Reproducible non-negative matrix factorization

NNMF decomposes the high-dimensional epoched EEG signal into several low rank approximations(Lee and Seung, 1999). Each rank is composed of two vectors, approximating the time domain (meta-MRPs, borrowing the language from genetics (Brunet et al., 2004)) or the corresponding prominence of each trial (*meta-Trials*, **Figure 1**). We applied non-negative matrix factorization to the signals surrounding each participants’ movements (−500 to 100 ms from the end of the thumb flexion) for the selected electrodes. The optimal number of ranks per participant was determined through cross-validation. Towards this, the epoched data was made non-negative by shifting it with the minimum voltage, and 30 percent of the data points were randomly removed (masking). To ensure stability in the solution of non-negative matrix factorization, which is a non-convex problem dependent on initialization, 100 repetitions with randomly initialized values was performed during the cross-validations (Wu et al., 2016). The meta-MRPs and meta-Trials were z-normalized and used to reconstructed the original trials x amplitude EEG tensor. With this reconstruction, the test error can be calculated. The best rank (between 2 and 10) was selected based on the smallest test error, averaged across all repetitions. Finally, the non-negative matrix factorization was repeated, with the best rank and without any masking, 1000 times with randomly initialized values. The most stable and reproducible result across the repetitions was selected, by first calculating the pairwise correlation coefficient between the decompositions of every repetition, and selecting the highest correlation. The median of the pairwise cross-correlation was calculated for every decomposition and the decomposition with the highest median across the 1000 repetitions was selected (reproducible NNMF method introduced in (Ceolini and Ghosh, 2022).

### Clustering non-negative matrix factorization decompositions

Patterns of meta-MRPs were identified across the sampled population with the k-means clustering algorithm (S. Lloyd, 1982). Resulted clusters could have meta-MRPs of the same participant but a different rank, as the meta-MRPs were considered independent during the procedure. The optimal number of clusters for k-means was selected based on the silhouette method considering between 2 and 8 clusters. This method selects the optimal number of clusters based on the squared Euclidean distance between the resulting clusters (Rousseeuw, 1987). The procedure was repeated 1000 times to account for the non-unique nature of K-means clustering (Lisboa et al., 2013). Five clusters were the most common across the repetitions. Finally, k-means with the selected number of clusters was repeated 1000 times and the repetition with the smallest sum of squared distance was selected as the most stable (Lisboa et al., 2013) (Figure 2). To verify the stability, we repeated the procedure for stable k-means 50 times and compared the clusters with cross-correlation. The final selected clusters were stable every repetition.

### Statistical analysis

The relationship between meta-MRPs and event-related potentials was analyzed by sorting trials based on meta-trial prominence. Pearson correlation coefficient was calculated between meta-MRPs and event-related potentials of the top 10 to 100 percent trials (incrementally increased by 10 percent). We then calculated the median correlation for each percentage and selected a threshold of 30 percent, derived from a trade-off with the median correlation (R = 0.4) values and the number of trials included (N = 131).

The event-related potentials were calculated with the top 30 percent of trials sorted by the meta-trials. They were analyzed using one-sample *t*-tests across all electrodes and time-points (−2 to 2 s). The event-related signals were estimated based on trimmed means (20%). The statistical corrections for multiple comparisons were performed through spatiotemporal clustering (α = 0.05, 1000 bootstraps) using the LIMO EEG toolbox (Pernet et al., 2011) **Figure 2, Figure 3**). The one-sample *t*-tests and multiple comparison corrections were also repeated on the population level with all the trials (Supplementary Figure 1).

#### Behavioral features extraction and pre-processing

To understand the relationship between the NNMF and behavioral features we trained, four regression models with the force, inter-event-intervals, duration, and movement onset. The force generated was the maximum voltage within 5 ms of the contact time, while the inter-event-interval was defined as the gap from the previous event. The duration was the distance between rise and fall of the signal. Finally, movement onset was identified when the variance of the signal abruptly changes (Killick et al., 2012). A robust regression model was fit (using bisquare weights) with each behavioral feature as independent variable and the meta-trials as dependent variable. Subsequently, we conducted Wilcoxen signrank tests with the beta coefficients of the participants in each cluster. Bonferroni correction (p < 0.05/4) was applied for multiple comparisons correction (Supplementary Figure 3).

### Data and code availability

Pending. The updated pre-print or publication will contain the code and shared data links.

## Acknowledgment

The authors would like to thank the student assistants who contributed to the data collection at Leiden University. The authors appreciate the discussions with Enea Ceolini.

## Related Figures

**Supplementary Figure 1**. Event-related potentials (blue line, trimmed means 20%) for selected sensorimotor electrodes of each participant. Single trials (black lines) demonstrate trial-by-trial variability. The black box shows the signal preparation period selected for non-negative matrix factorization

**Supplementary Figure 2**. Grand average event-related potentials of EEG signal surrounding force sensor contact. **(a)** Negative pre-movement processing is visible at the left sensorimotor electrodes contralateral to the touch (left plot). The event-related potential is shown with trimmed means (20%) and 95% confidence intervals, significant timepoints are shaded purple (MCC, p < 0.05). Post-movement positivity is clearer at central electrode (right plot). **(b)** Scalp topologies of trimmed means (20%) and T-values show significant bilateral to contralateral activity preceding the event. Followed by positive sensory processing.

**Supplementary Figure 6**. Number of trials per participant. An average of (434) trials remained after pre-processing (red) as opposed to (553) collected trials (dark red).

## Related Figures

**Supplementary Figure 5**. Paired samples *t*-tests between cluster a and b show statistically significant differences in neural processing between the clusters (MCC, p < 0.05). Also, see Supplementary Movie x for the full statistical outcomes of the paired *t*-test.

**Supplementary Figure 3**. The relationship between the meta-trial and subsequent meta-trial scatter plot for all participants.

**Supplementary Figure 4**. Empirical cumulative distributions of beta-coefficients per behavioral feature show no clear relationship in any cluster.

## Movies

**Supplementary Movie 1**. Grand average event-related potentials surrounding force sensor contact. T-values based on one sample *t*-tests corrected for multiple comparisons (MCC, p < 0.05).

**Supplementary Movie 2**. Event-related potentials and T-values surrounding force sensor contact for participants in first cluster (cluster a in Figure 2 and 3).

**Supplementary Movie 3**. Event-related potentials and T-values surrounding force sensor contact for participants in second cluster (cluster b in Figure 2 and 3).

**Supplementary Movie 4**. Event-related potentials and T-values surrounding force sensor contact for participants in third cluster (cluster c in Figure 2 and 3).

**Supplementary Movie 5**. Event-related potentials and T-values surrounding force sensor contact for participants in fourth cluster (cluster d in Figure 2 and 3).

**Supplementary Movie 6**. Event-related potentials and T-values surrounding force sensor contact for participants in fifth cluster (cluster e in Figure 2 and 3).

**Supplementary Movie 7**. Paired-sample *t*-test comparing event-related potentials of the first (a in Figure 2 and 3) and second (b in Figure 2 and 3) cluster. Statistics corrected for multiple comparisons (MCC, p < 0.05).

