## Supplementary Figure 1 for "Multiple forms of neural processing when repeating voluntary thumb flexions"

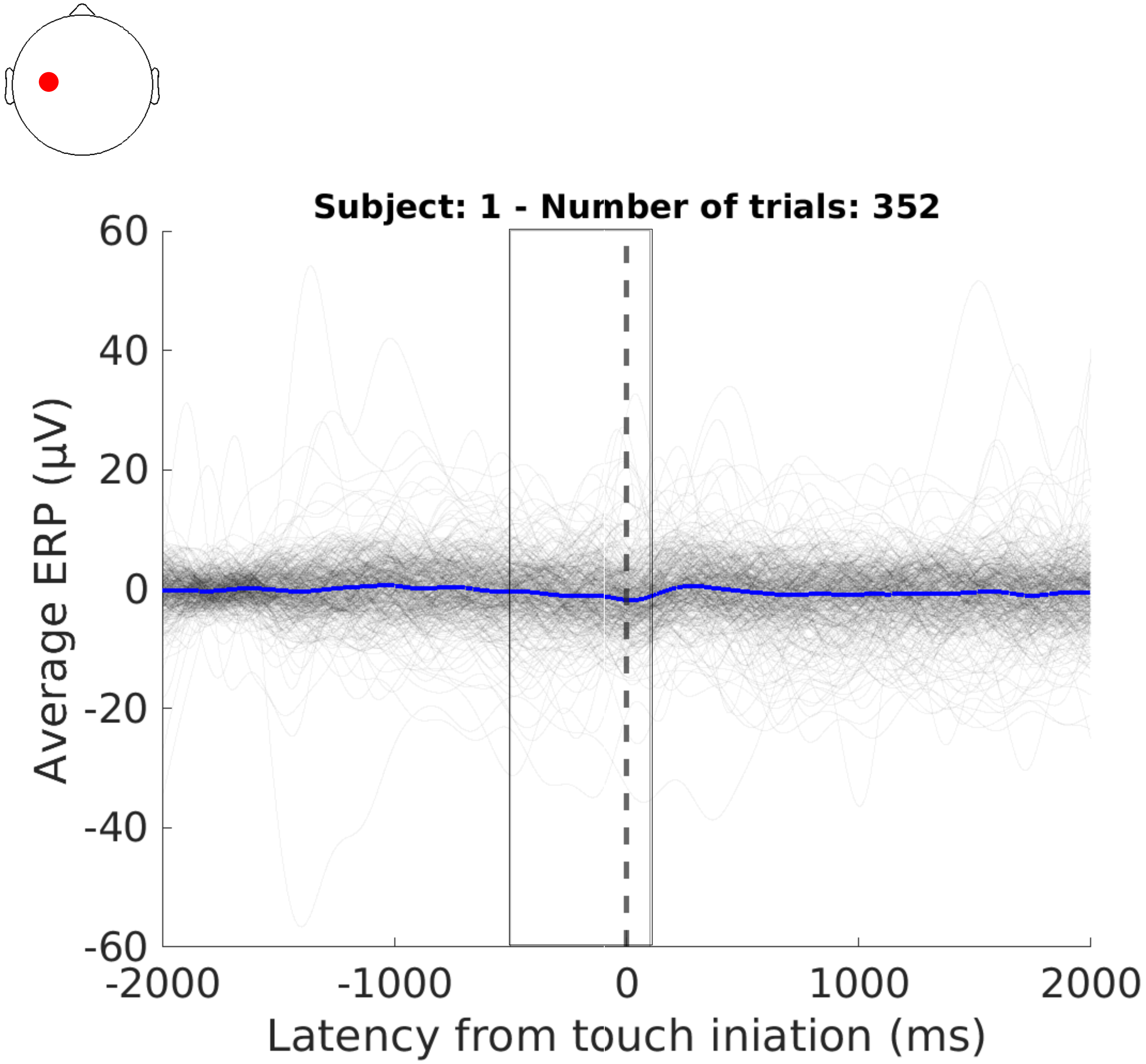
Subject: 1 - Number of trials: 352
Average ERP (µV)
60
40
20
0
-20
-40
-60
-2000
-1000
0
1000
2000
Latency from touch iniation (ms)

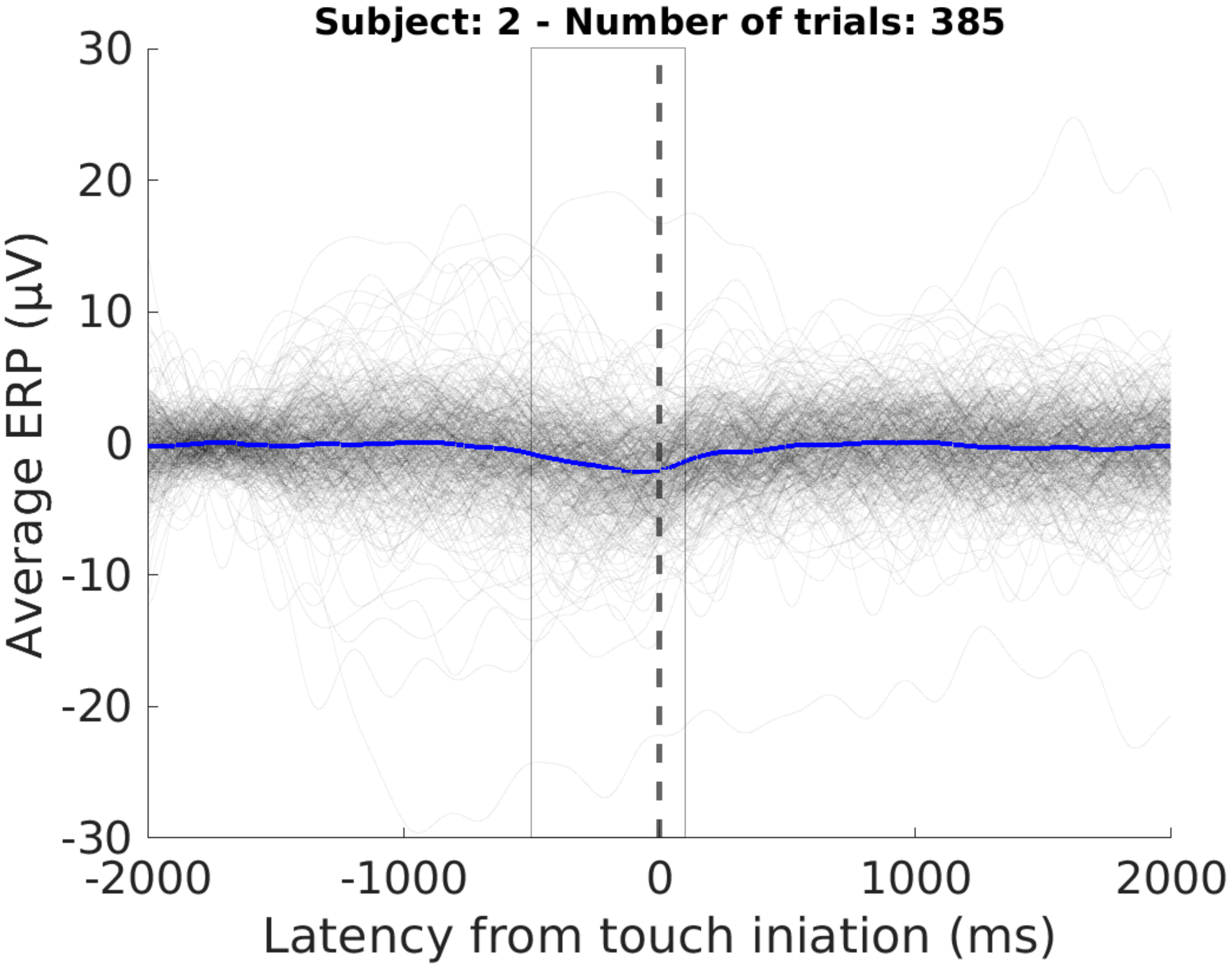
Subject: 2 - Number of trials: 385
30
20
10
0
-10
-20
-30
Average ERP (μV)
-2000
-1000
0
1000
2000
Latency from touch iniation (ms)

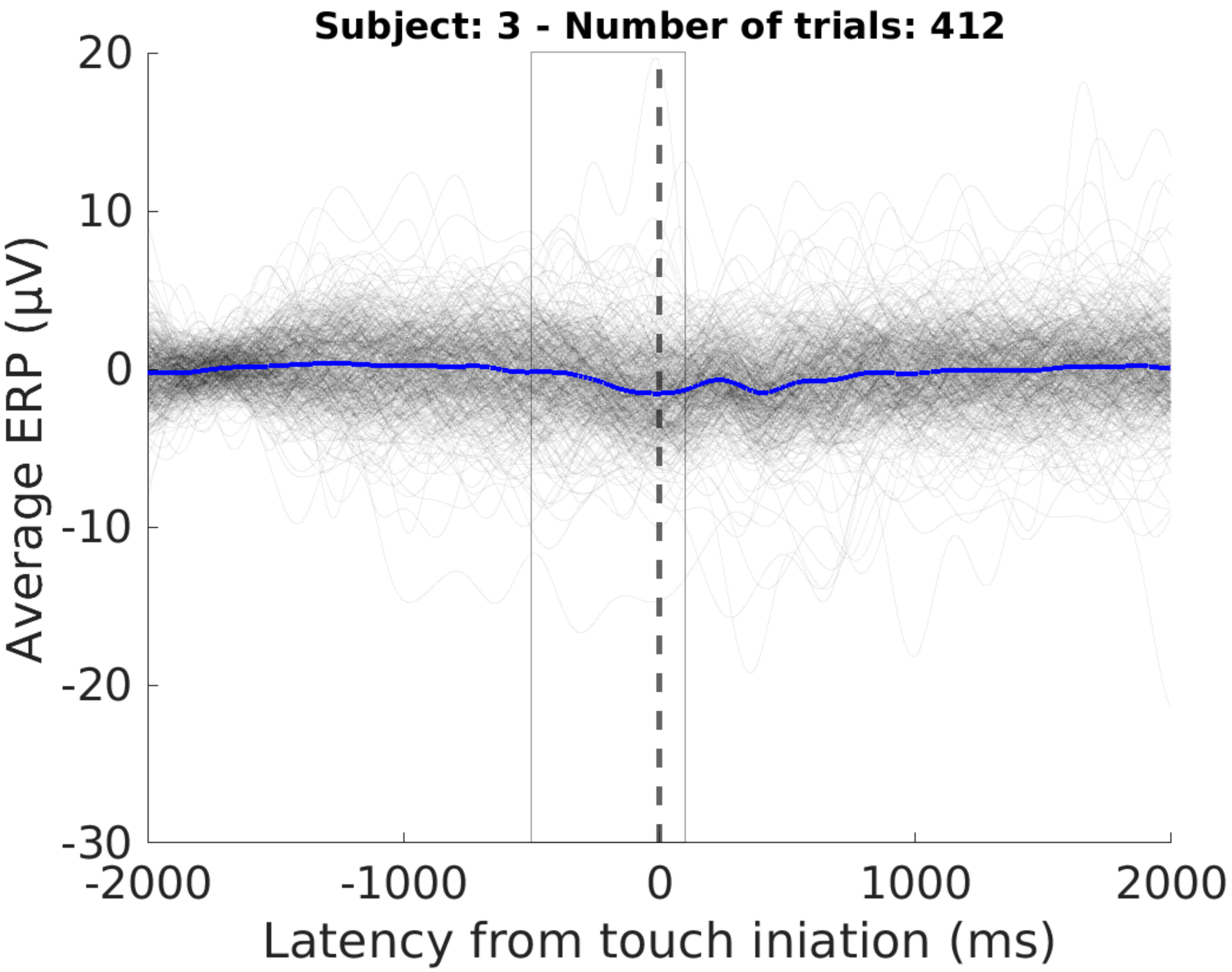
Subject: 3 - Number of trials: 412
Average ERP (µV)
20
10
0
-10
-20
-30
-2000
-1000
0
1000
2000
Latency from touch iniation (ms)

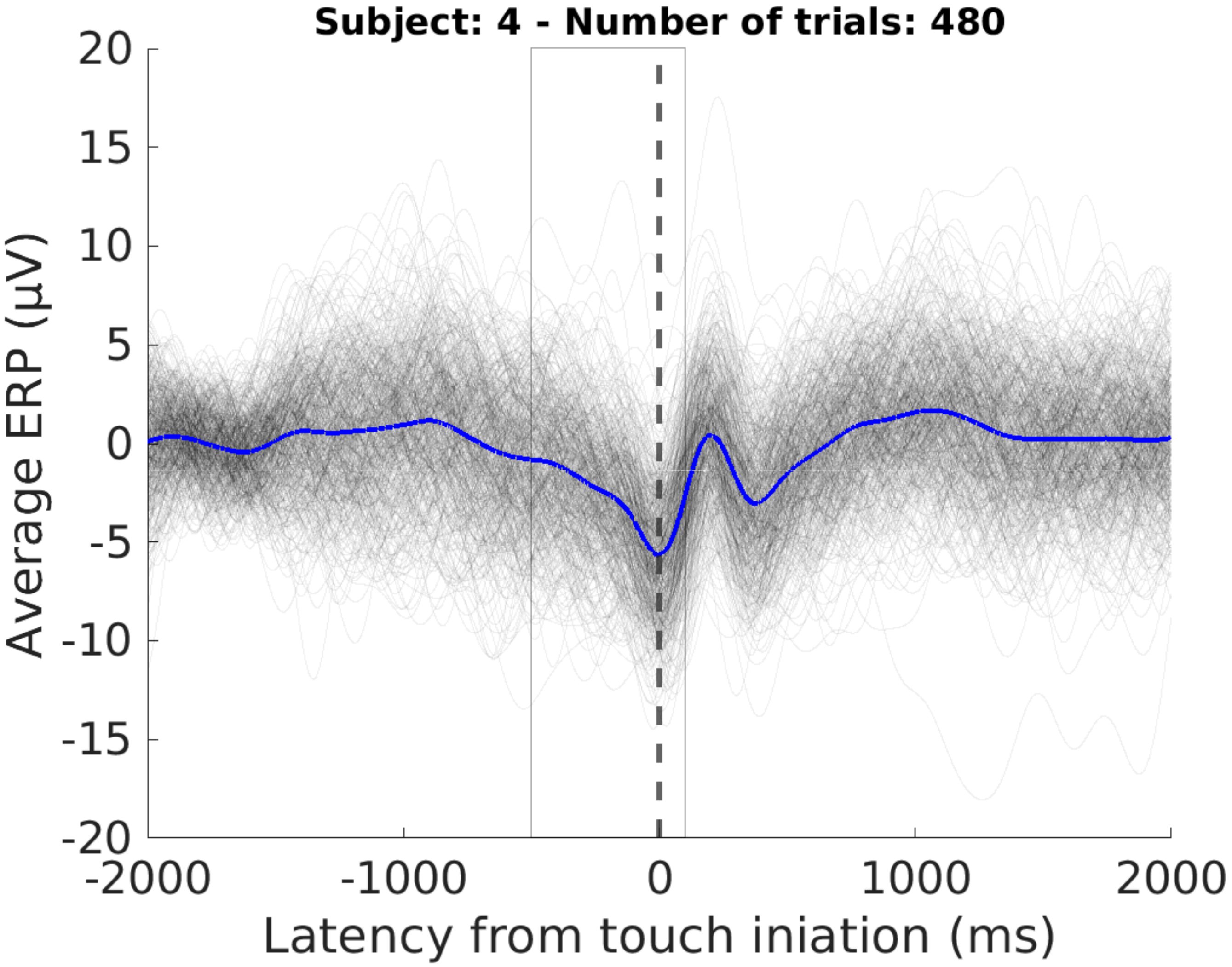

Subject: 4 - Number of trials: 480
20
15
10
5
0
-5
-10
-15
-20
Average ERP (µV)
-2000
-1000
0
1000
2000
Latency from touch iniation (ms)

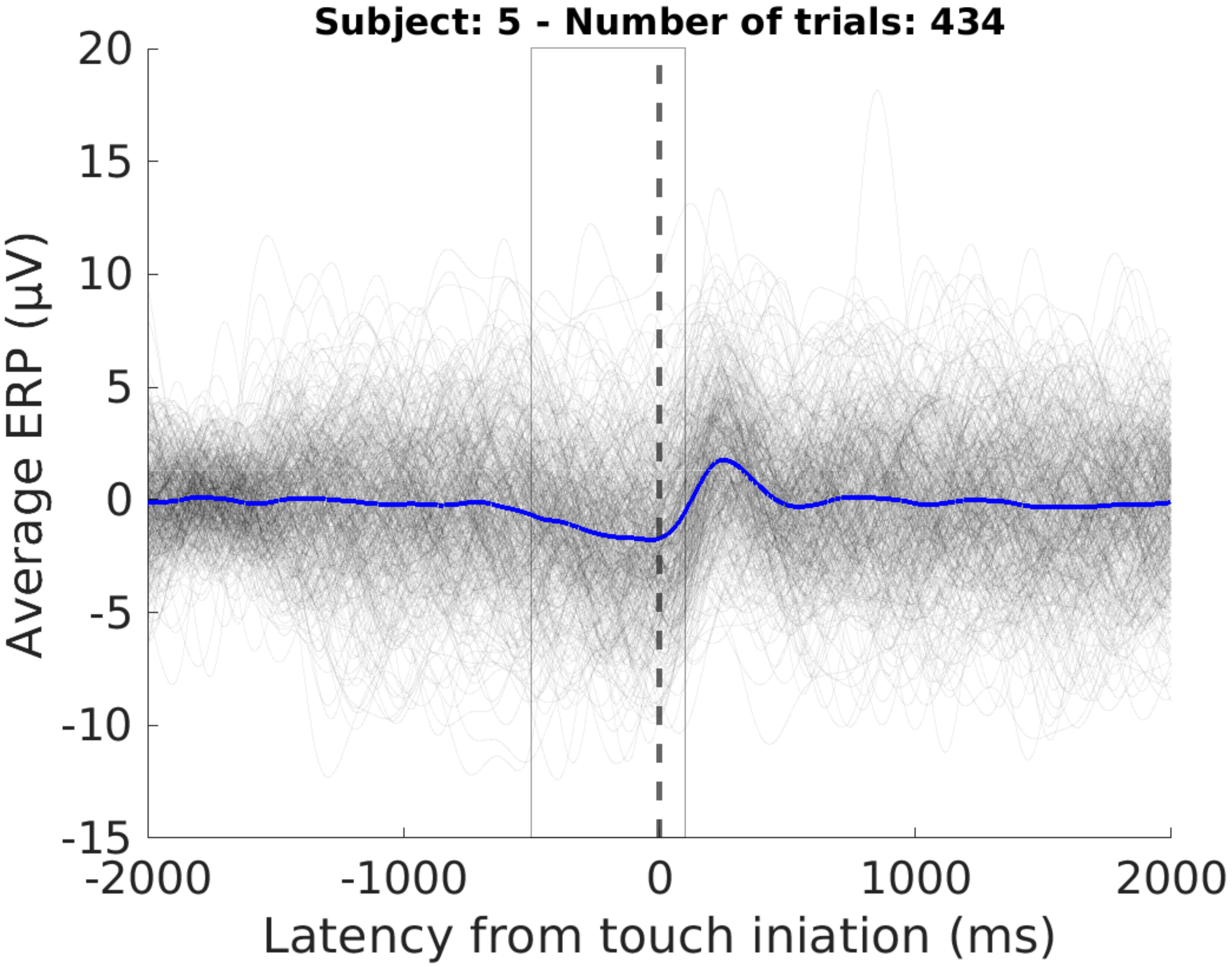

Subject: 5 - Number of trials: 434
20
15
10
5
0
-5
-10
-15
Average ERP (µV)
-2000
-1000
0
1000
2000
Latency from touch iniation (ms)

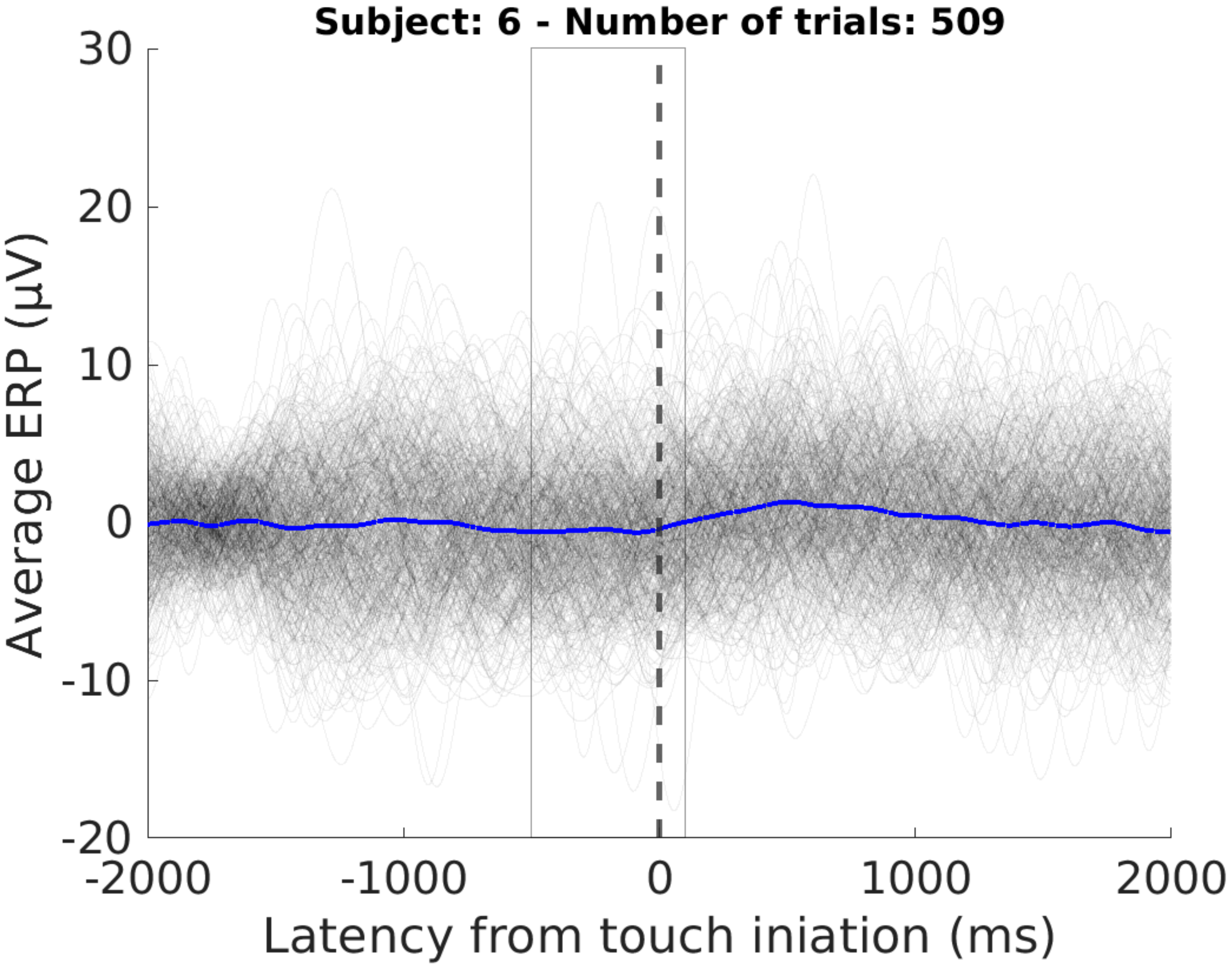

Subject: 6 - Number of trials: 509
30
20
10
0
-10
-20
Average ERP (μV)
-2000
-1000
0
1000
2000
Latency from touch iniation (ms)

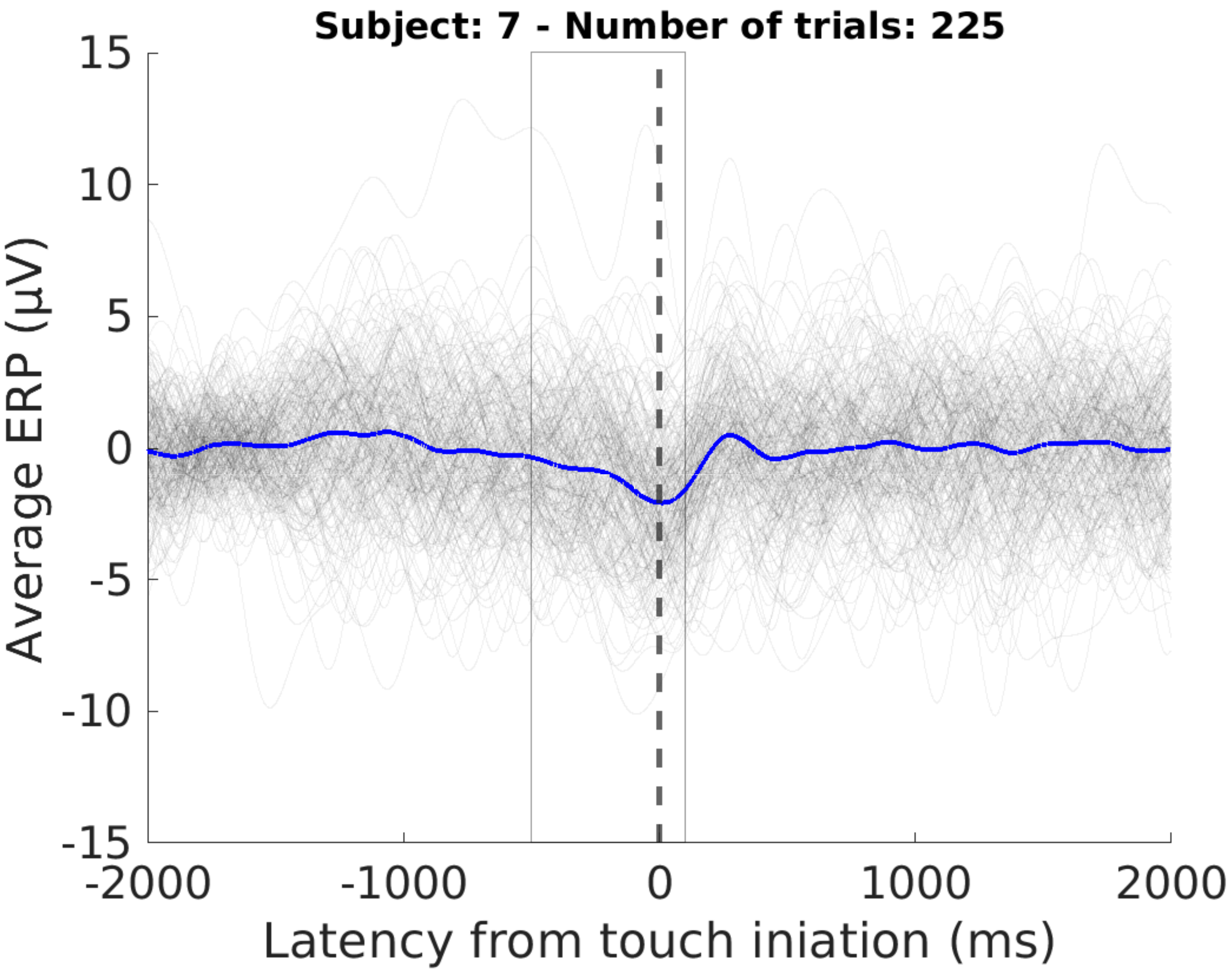
Subject: 7 - Number of trials: 225
15
10
5
0
-5
-10
-15
Average ERP (µV)
-2000
-1000
0
1000
2000
Latency from touch iniation (ms)

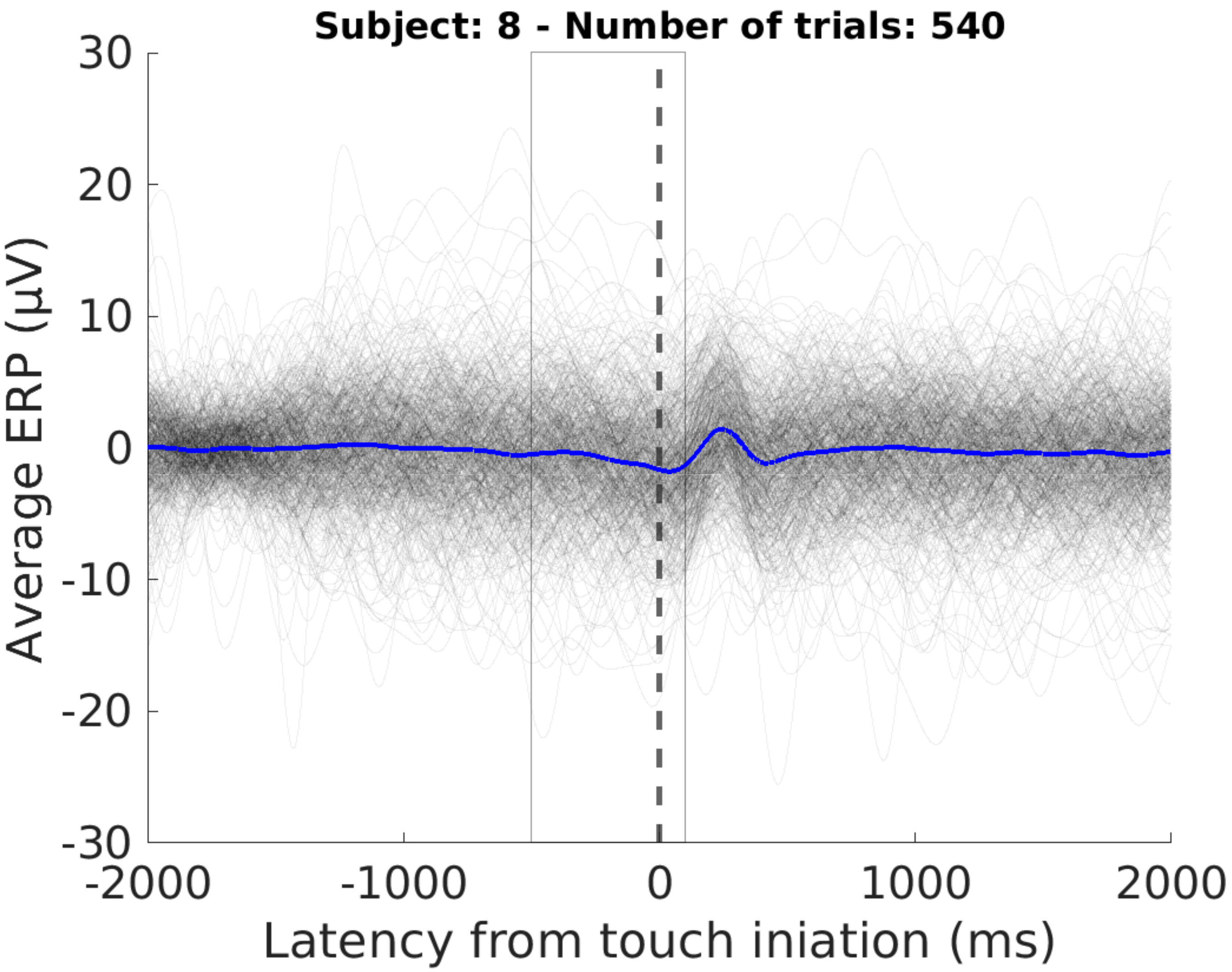

Subject: 8 - Number of trials: 540
30
20
10
0
-10
-20
-30
Average ERP (μV)
-2000
-1000
0
1000
2000
Latency from touch iniation (ms)

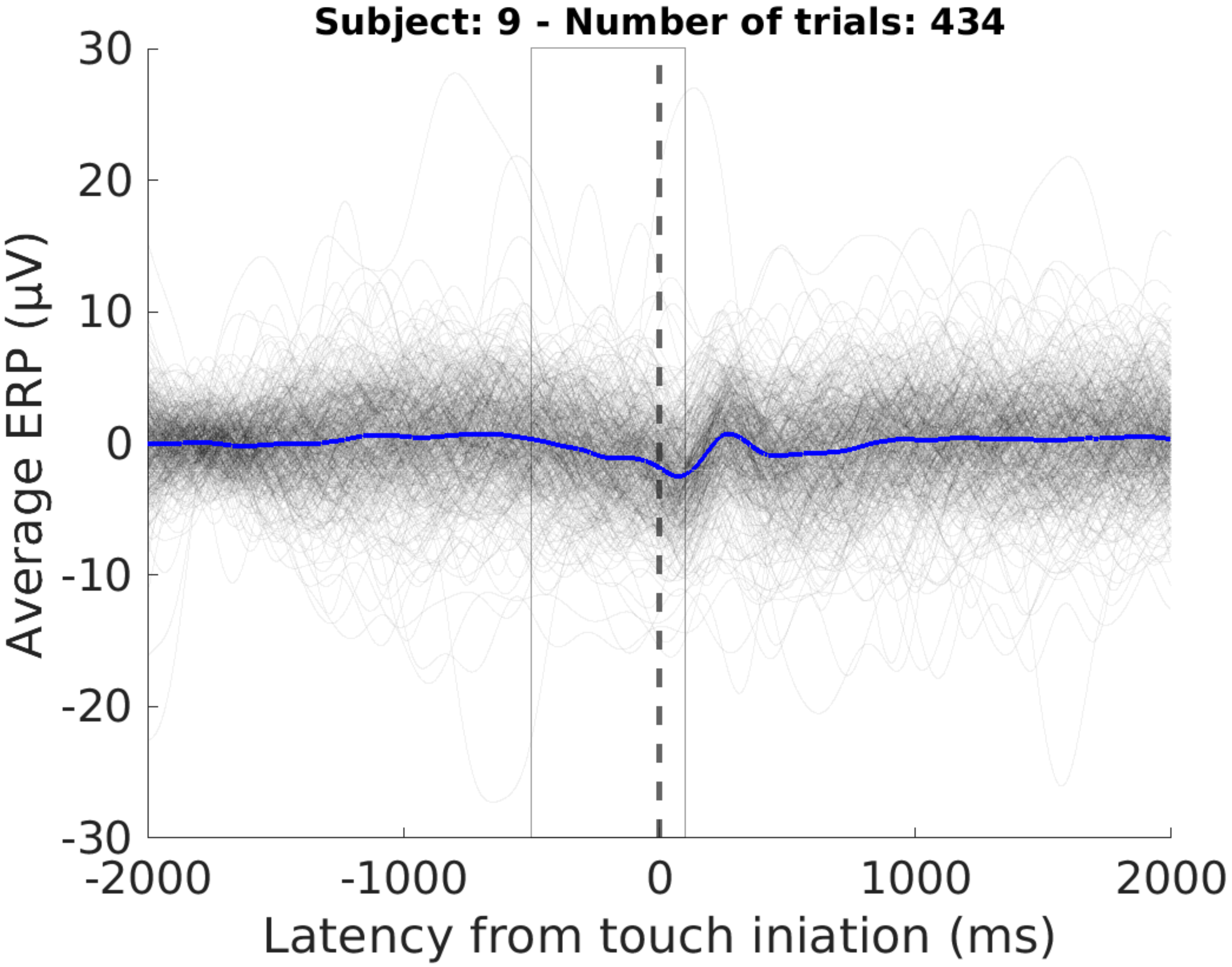
Subject: 9 - Number of trials: 434
30
20
10
0
-10
-20
-30
Average ERP (µV)
-2000
-1000
0
1000
2000
Latency from touch iniation (ms)

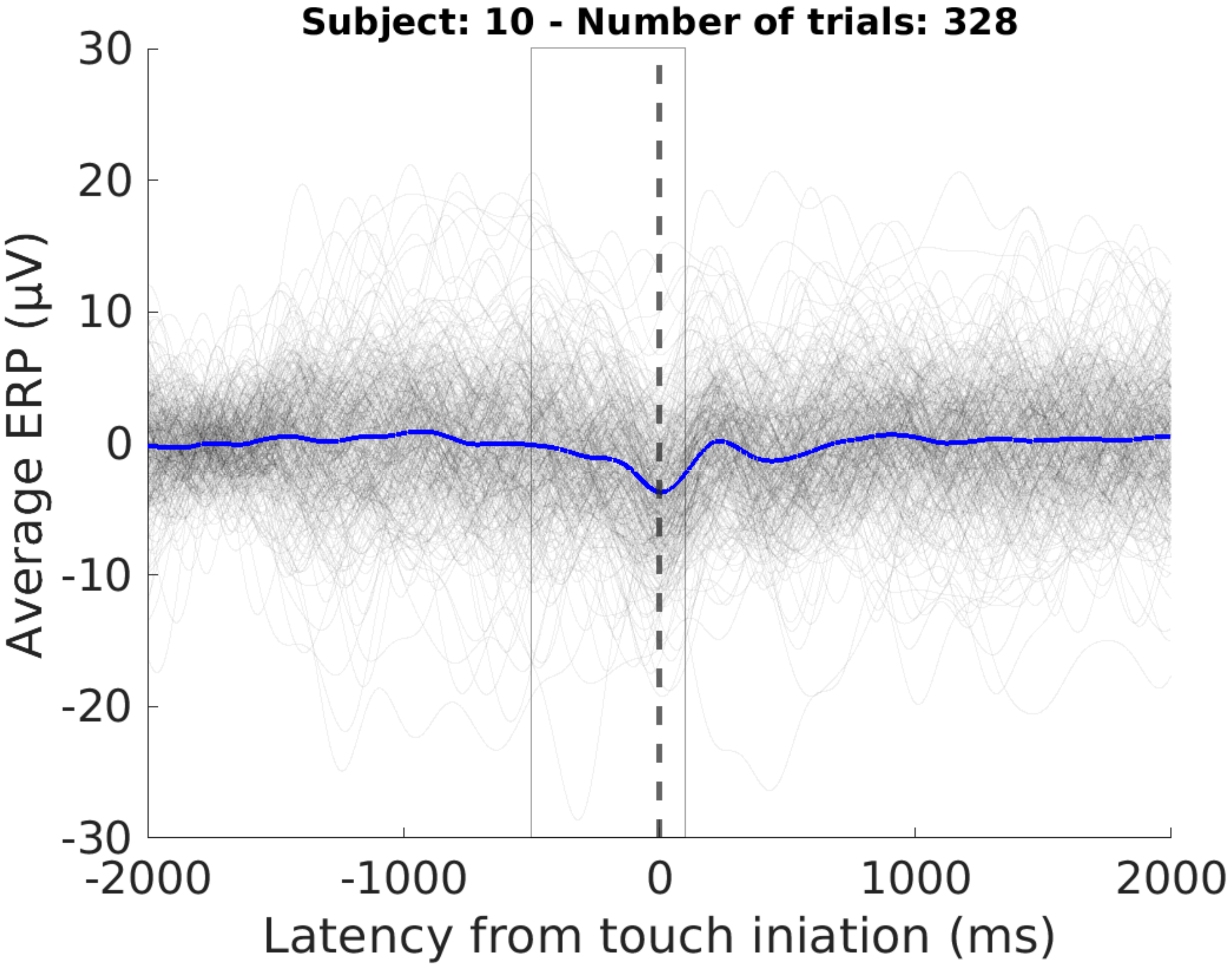

Subject: 10 - Number of trials: 328
30
20
10
0
-10
-20
-30
Average ERP (µV)
-2000
-1000
0
1000
2000
Latency from touch iniation (ms)

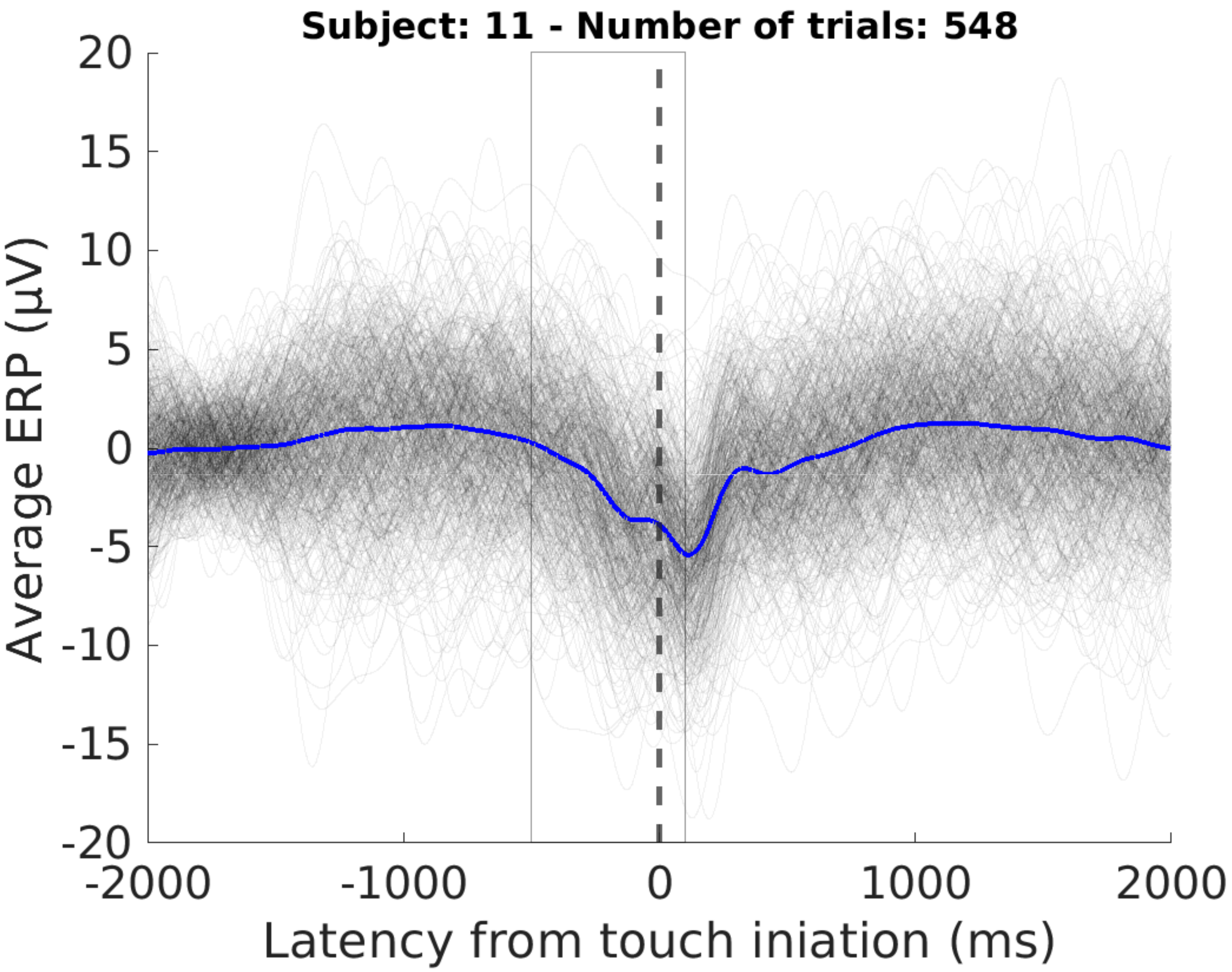

Subject: 11 - Number of trials: 548
Average ERP (µV)
20
15
10
5
0
-5
-10
-15
-20
-2000
-1000
0
1000
2000
Latency from touch iniation (ms)

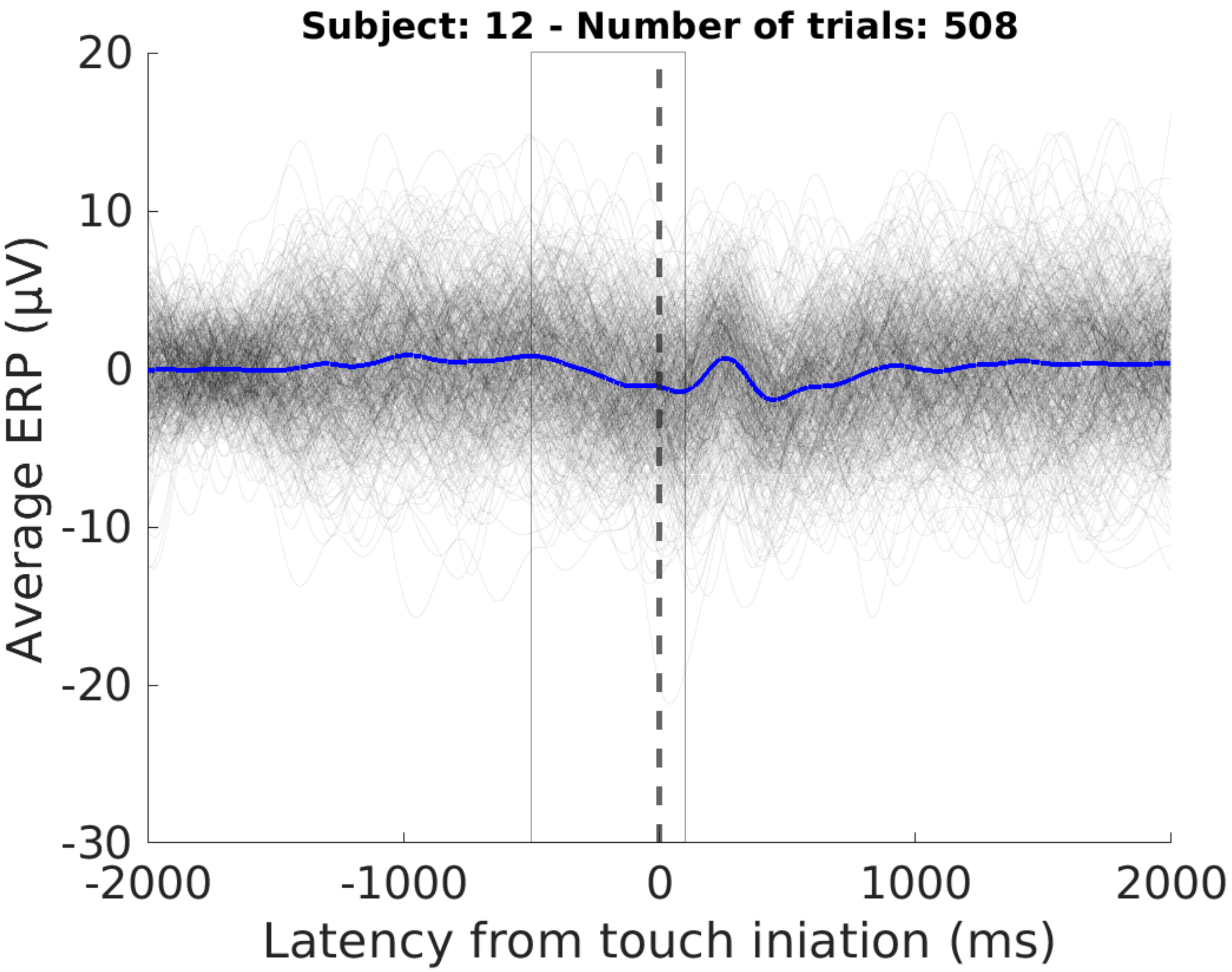
Subject: 12 - Number of trials: 508
20
10
0
-10
-20
-30
Average ERP (μV)
-2000
-1000
0
1000
2000
Latency from touch iniation (ms)

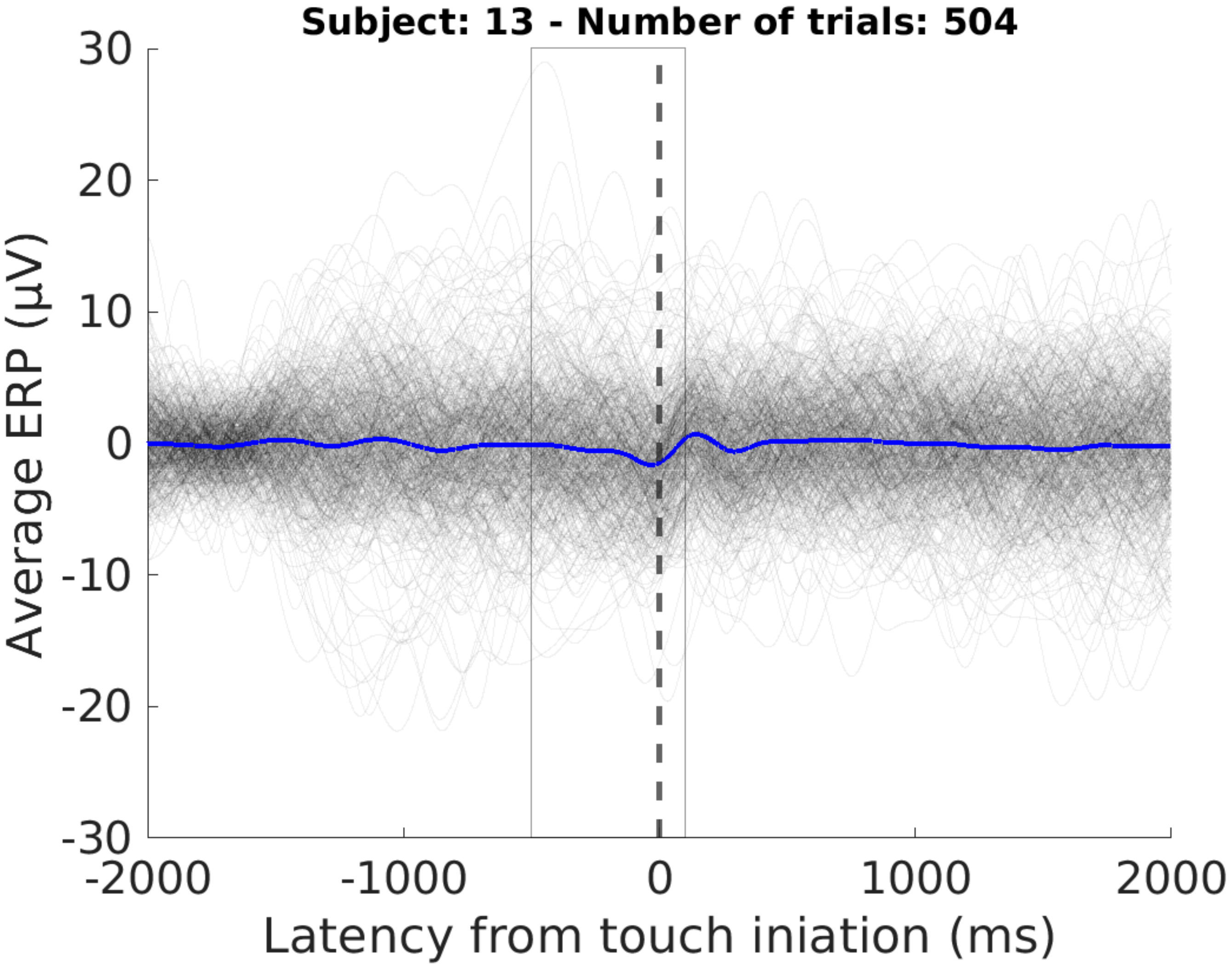

Subject: 13 - Number of trials: 504
30
20
10
0
-10
-20
-30
Average ERP (µV)
-2000
-1000
0
1000
2000
Latency from touch iniation (ms)

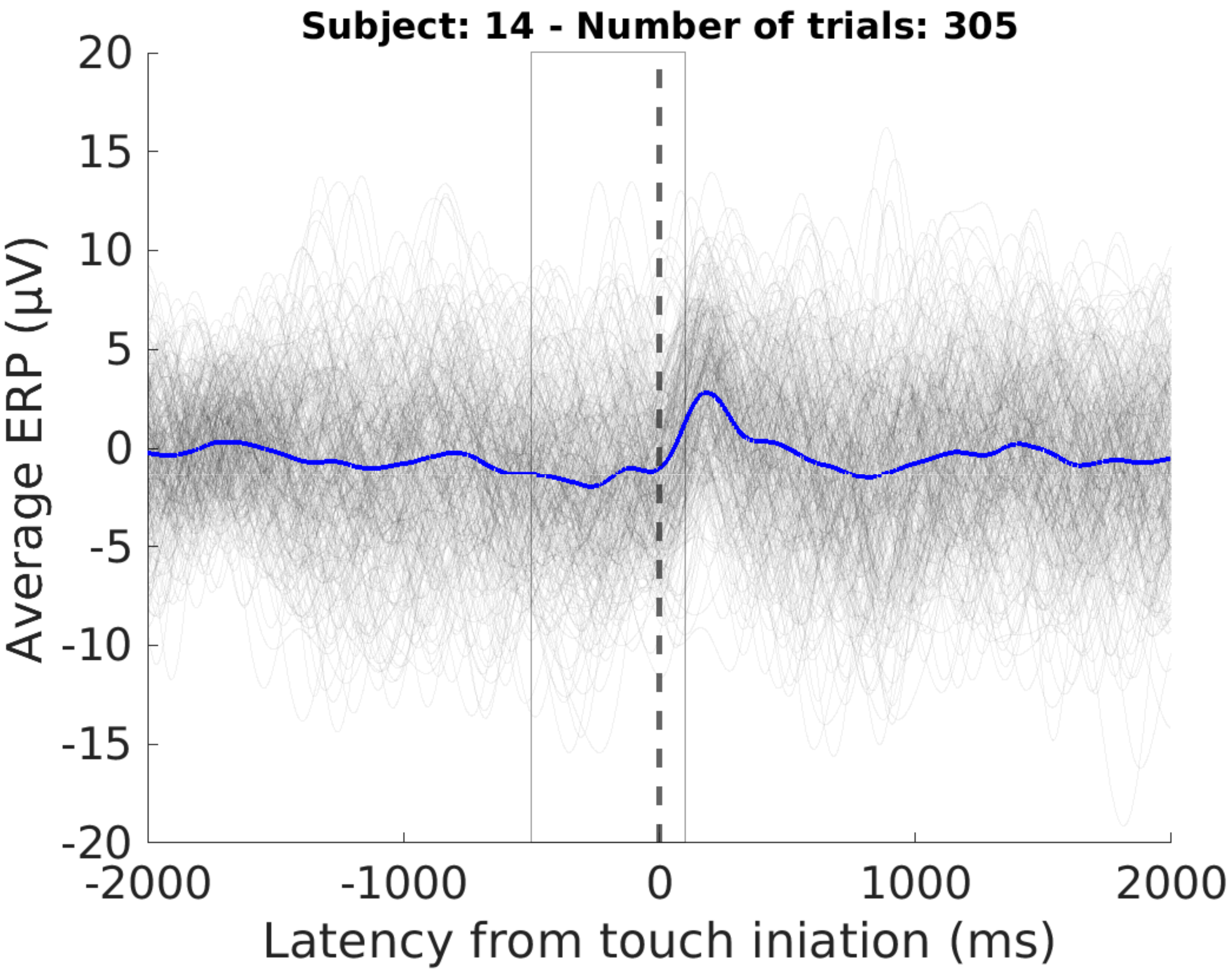

Subject: 14 - Number of trials: 305
Average ERP (μV)
20
15
10
5
0
-5
-10
-15
-20
-2000
-1000
0
1000
2000
Latency from touch iniation (ms)

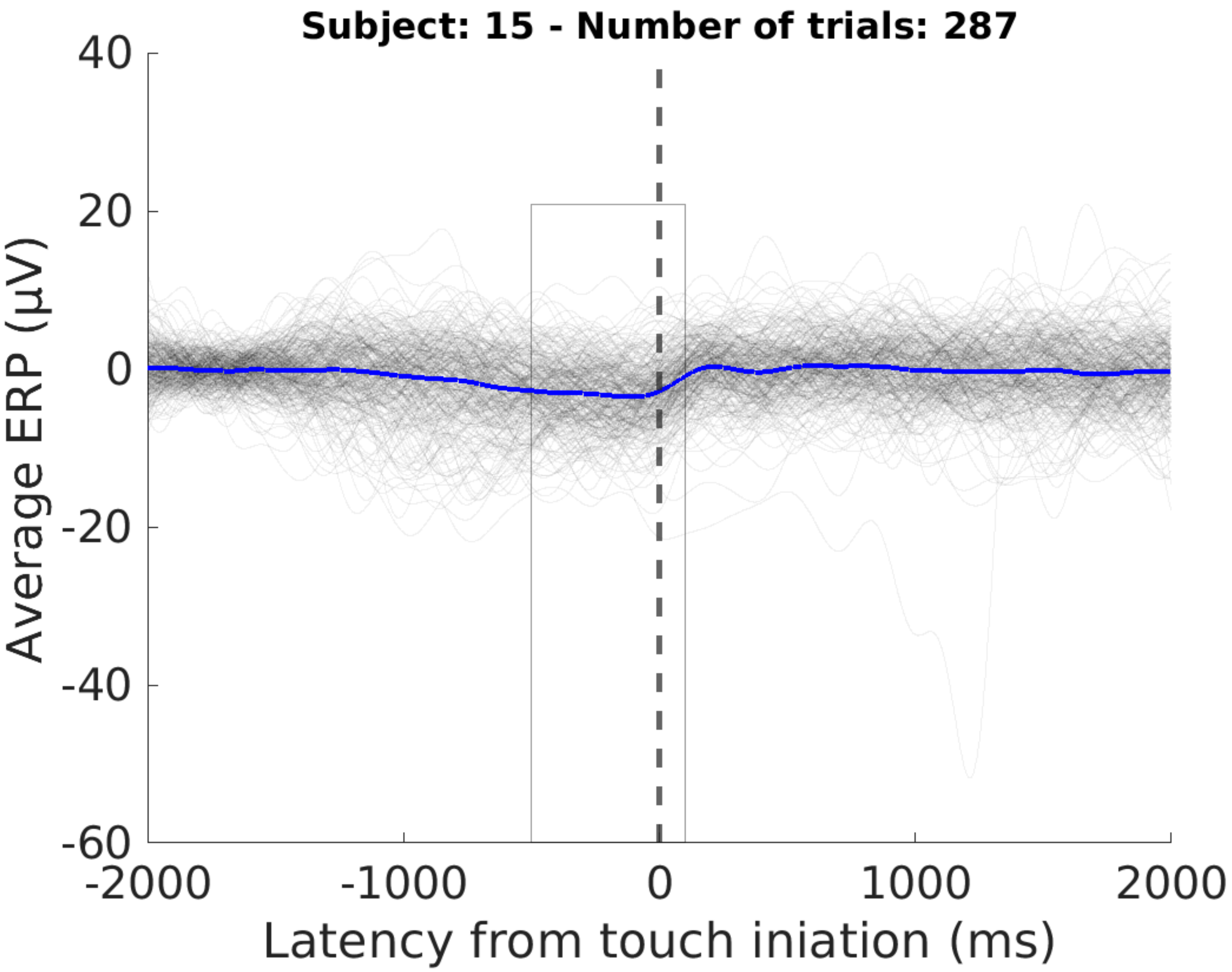

Subject: 15 - Number of trials: 287
40
20
0
-20
-40
-60
Average ERP (µV)
-2000
-1000
0
1000
2000
Latency from touch iniation (ms)

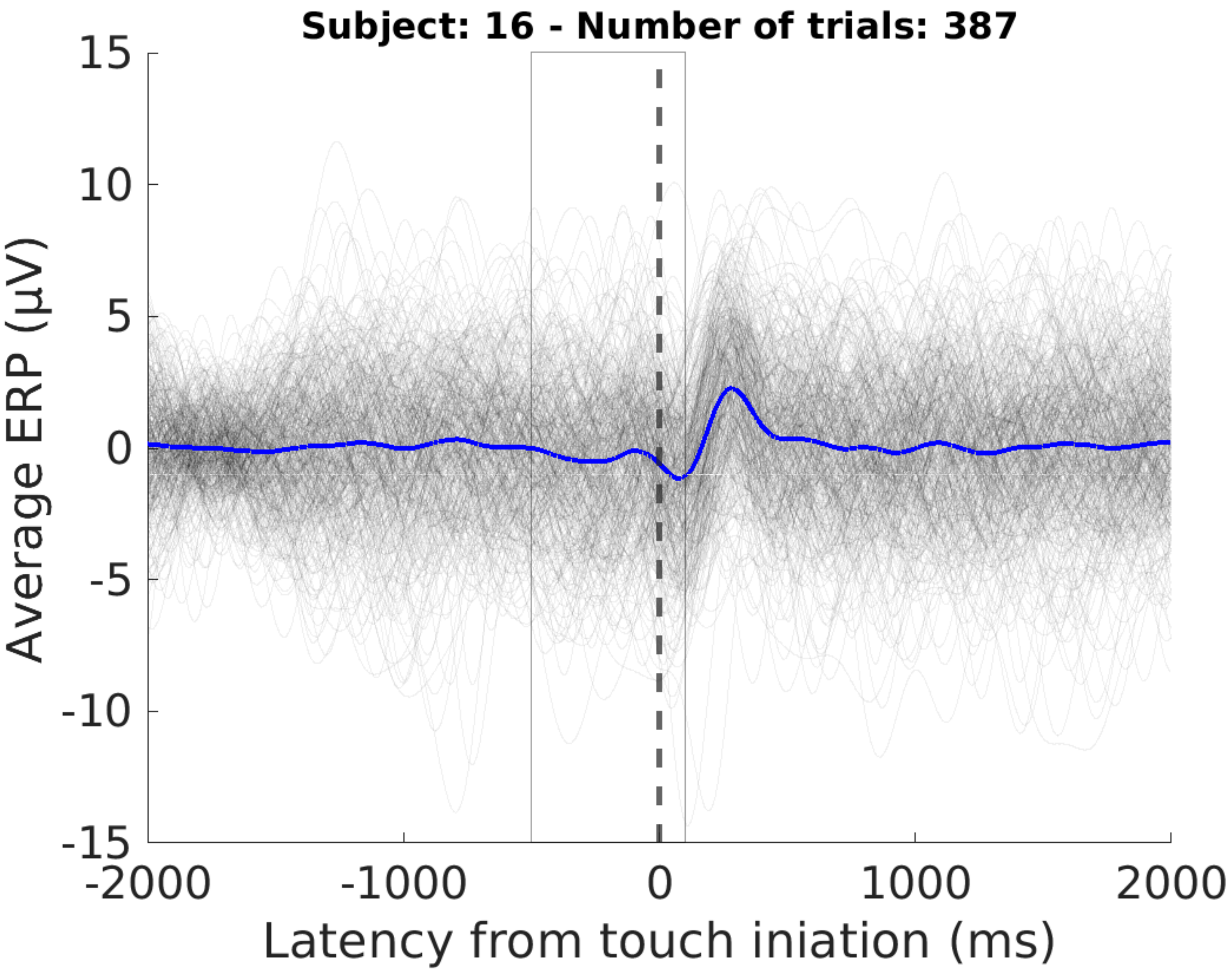
Subject: 16 - Number of trials: 387
15
10
5
0
-5
-10
-15
Average ERP (µV)
-2000
-1000
0
1000
2000
Latency from touch iniation (ms)

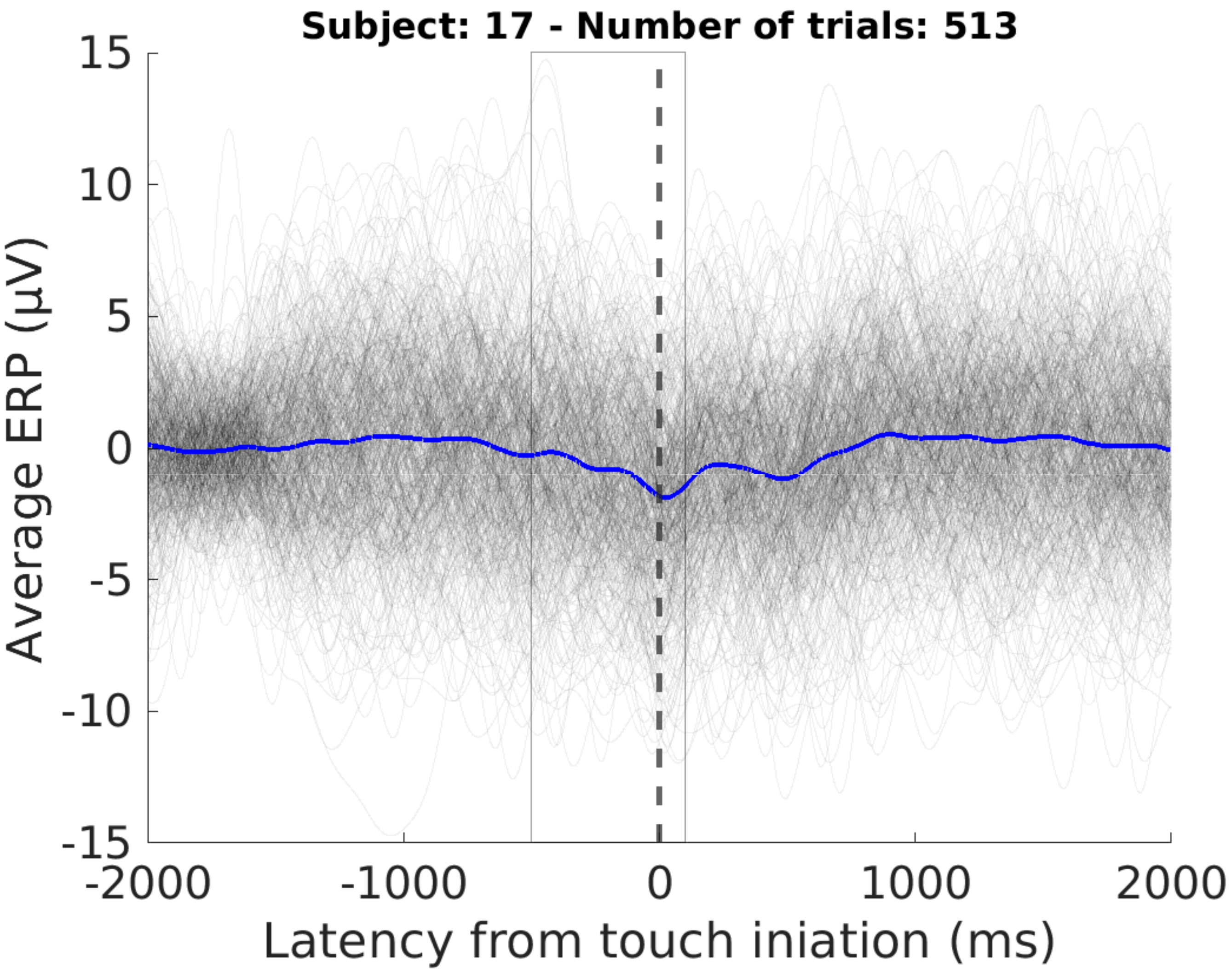
Subject: 17 - Number of trials: 513
15
10
5
0
-5
-10
-15
Average ERP (µV)
-2000
-1000
0
1000
2000
Latency from touch iniation (ms)

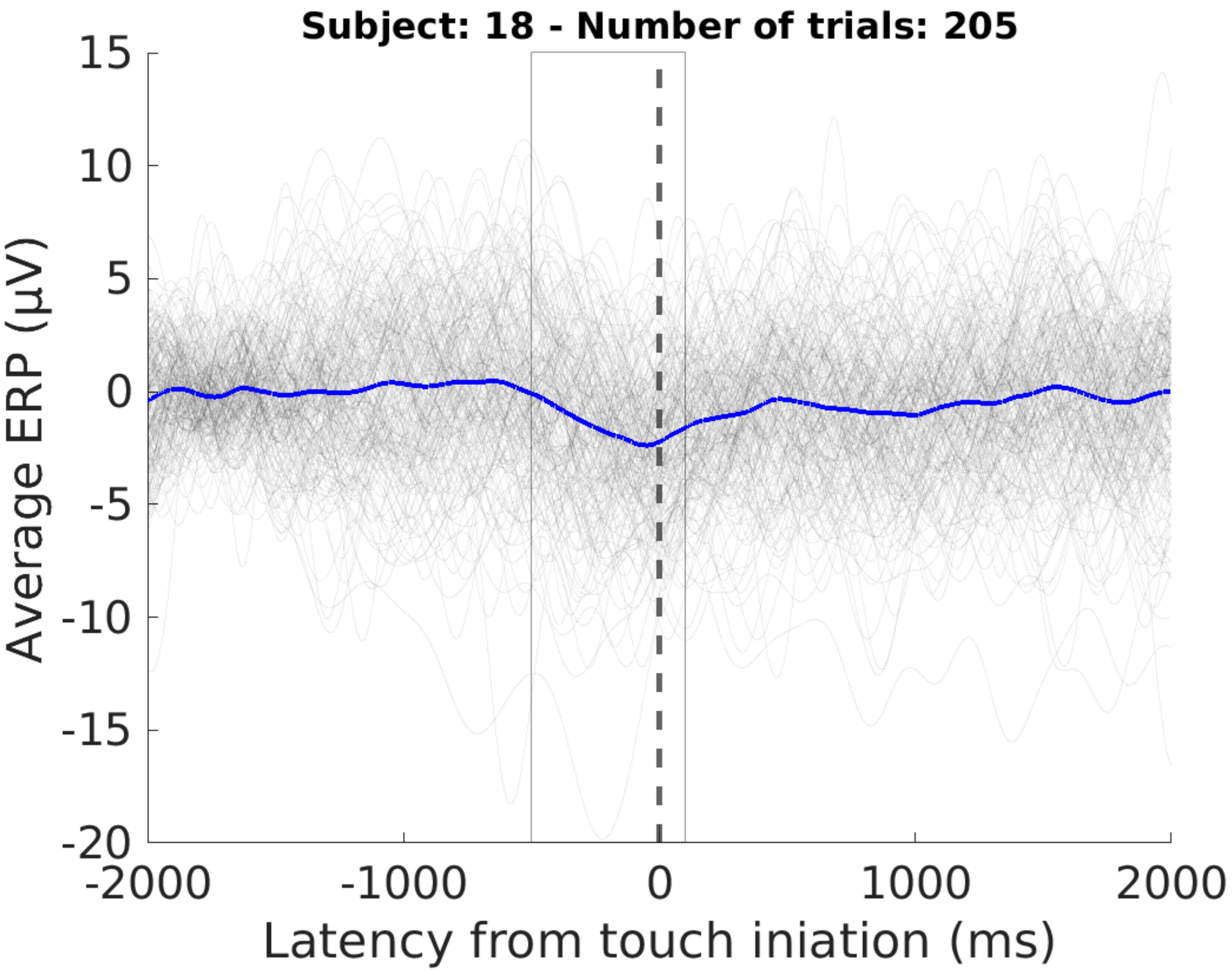

Subject: 18 - Number of trials: 205
15
10
5
0
-5
-10
-15
-20
Average ERP (µV)
-2000
-1000
0
1000
2000
Latency from touch iniation (ms)

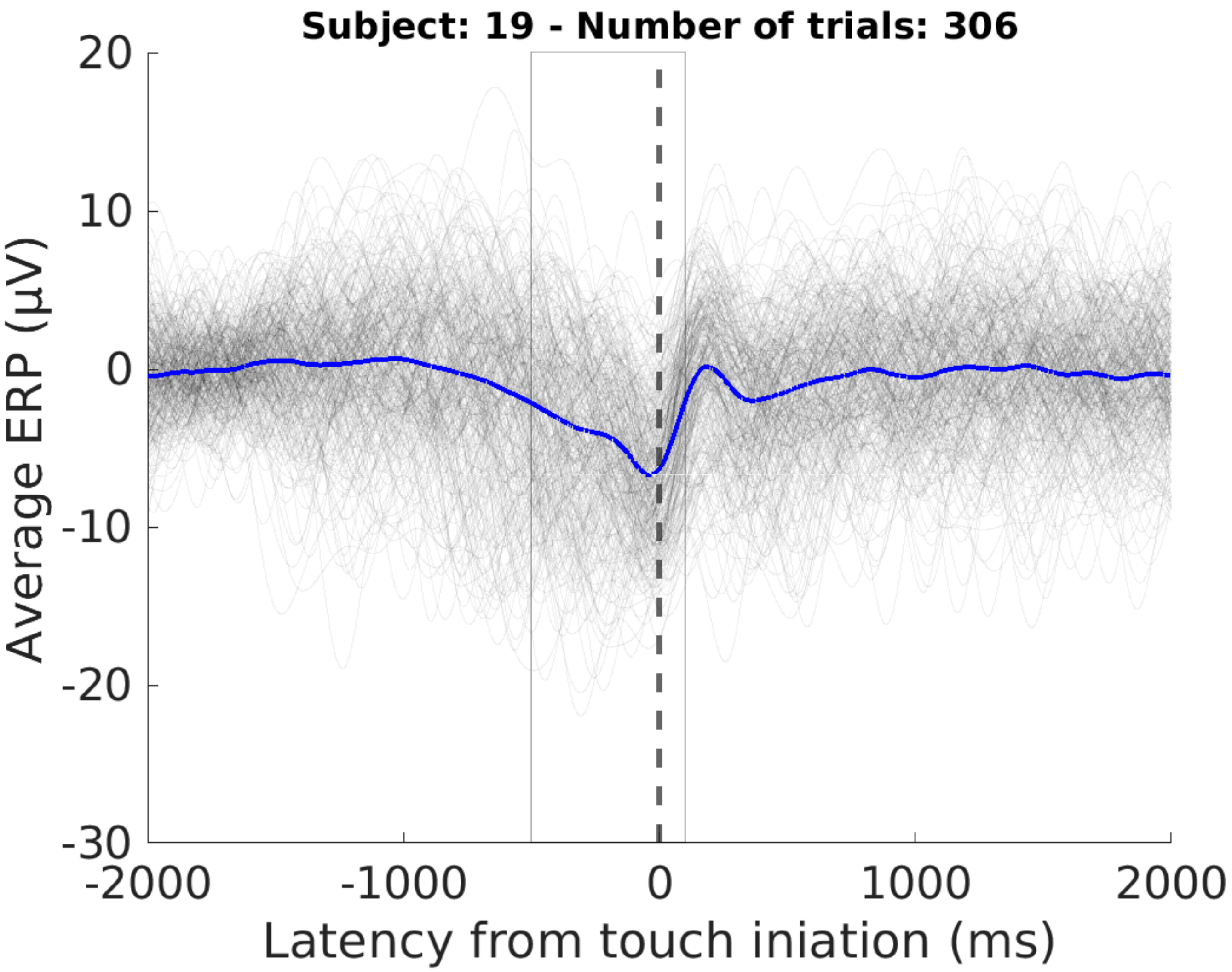
Subject: 19 - Number of trials: 306
20
10
0
-10
-20
-30
Average ERP (µV)
-2000
-1000
0
1000
2000
Latency from touch iniation (ms)

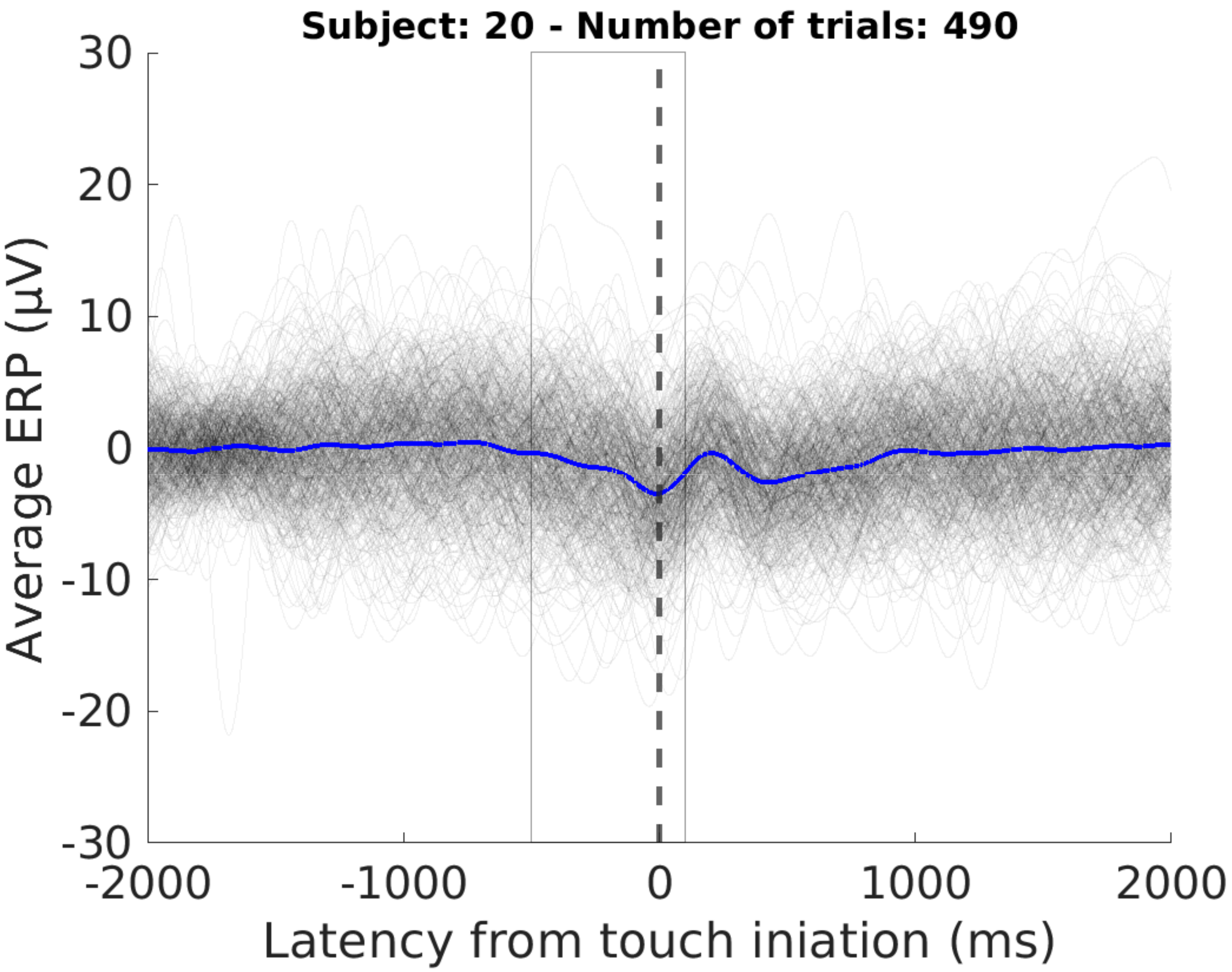
Subject: 20 - Number of trials: 490
Average ERP (µV)
30
20
10
0
-10
-20
-30
-2000
-1000
0
1000
2000
Latency from touch iniation (ms)

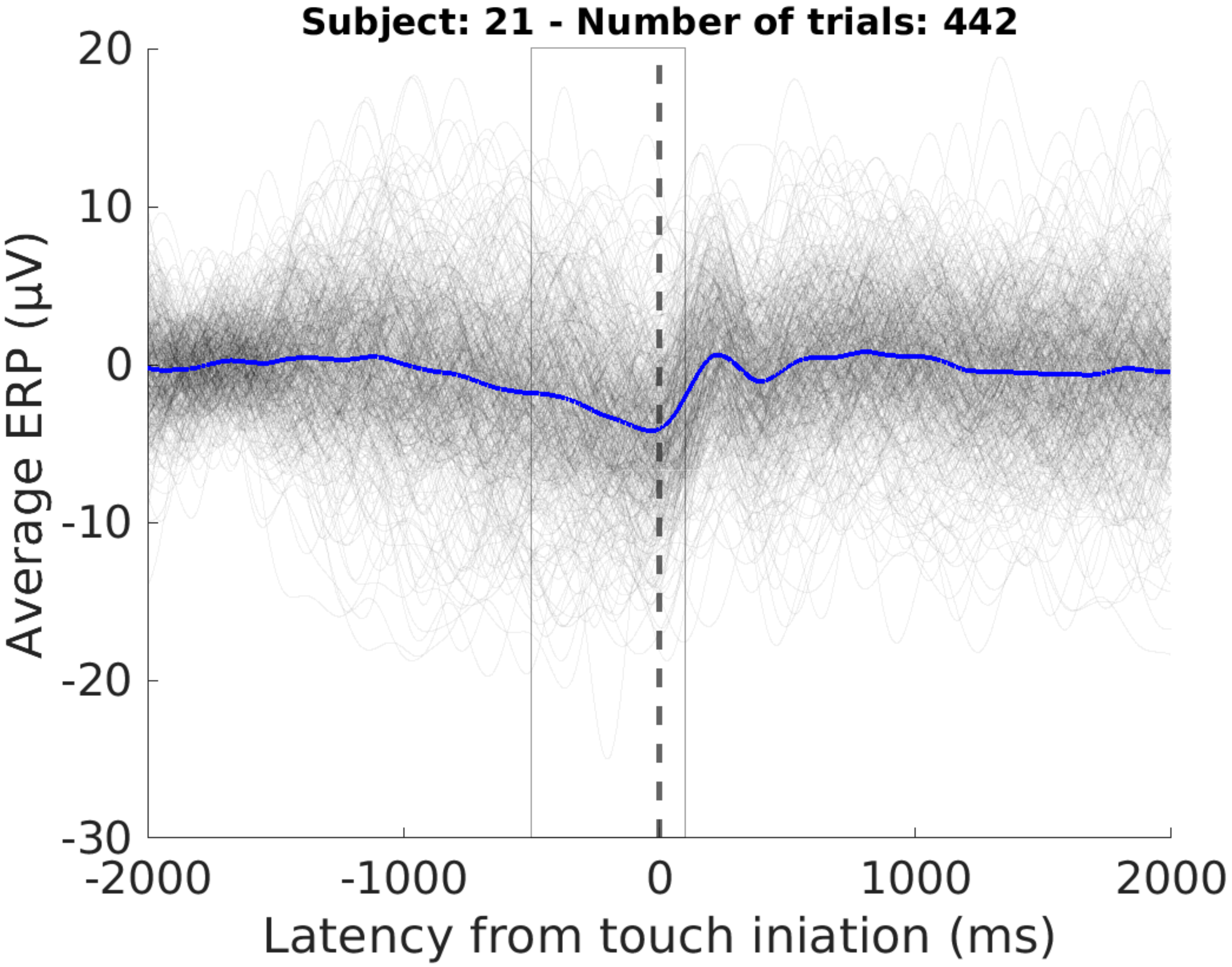
Subject: 21 - Number of trials: 442
20
10
0
-10
-20
-30
Average ERP (μV)
-2000
-1000
0
1000
2000
Latency from touch iniation (ms)

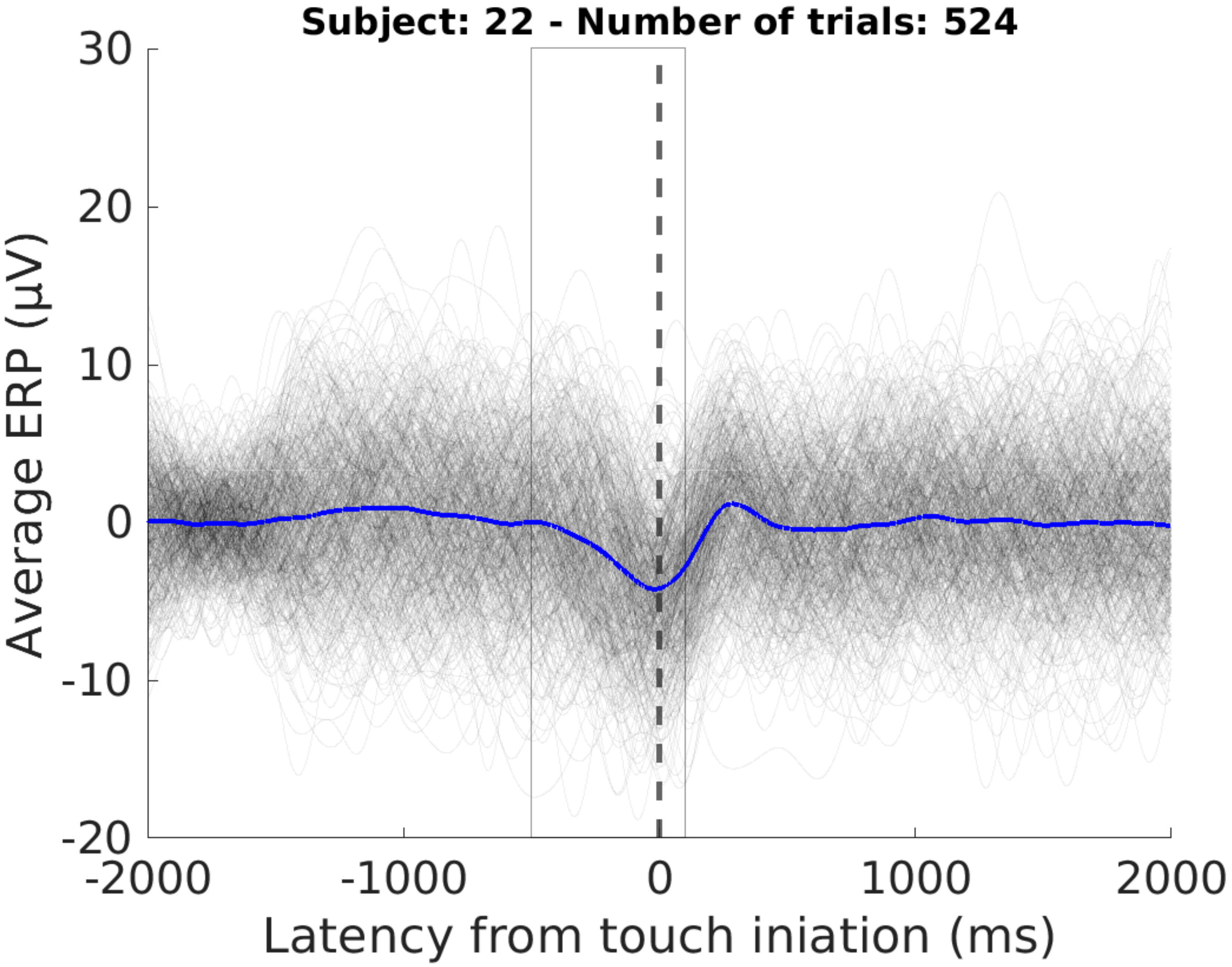
Subject: 22 - Number of trials: 524
30
20
10
0
-10
-20
Average ERP (μV)
-2000
-1000
0
1000
2000
Latency from touch iniation (ms)

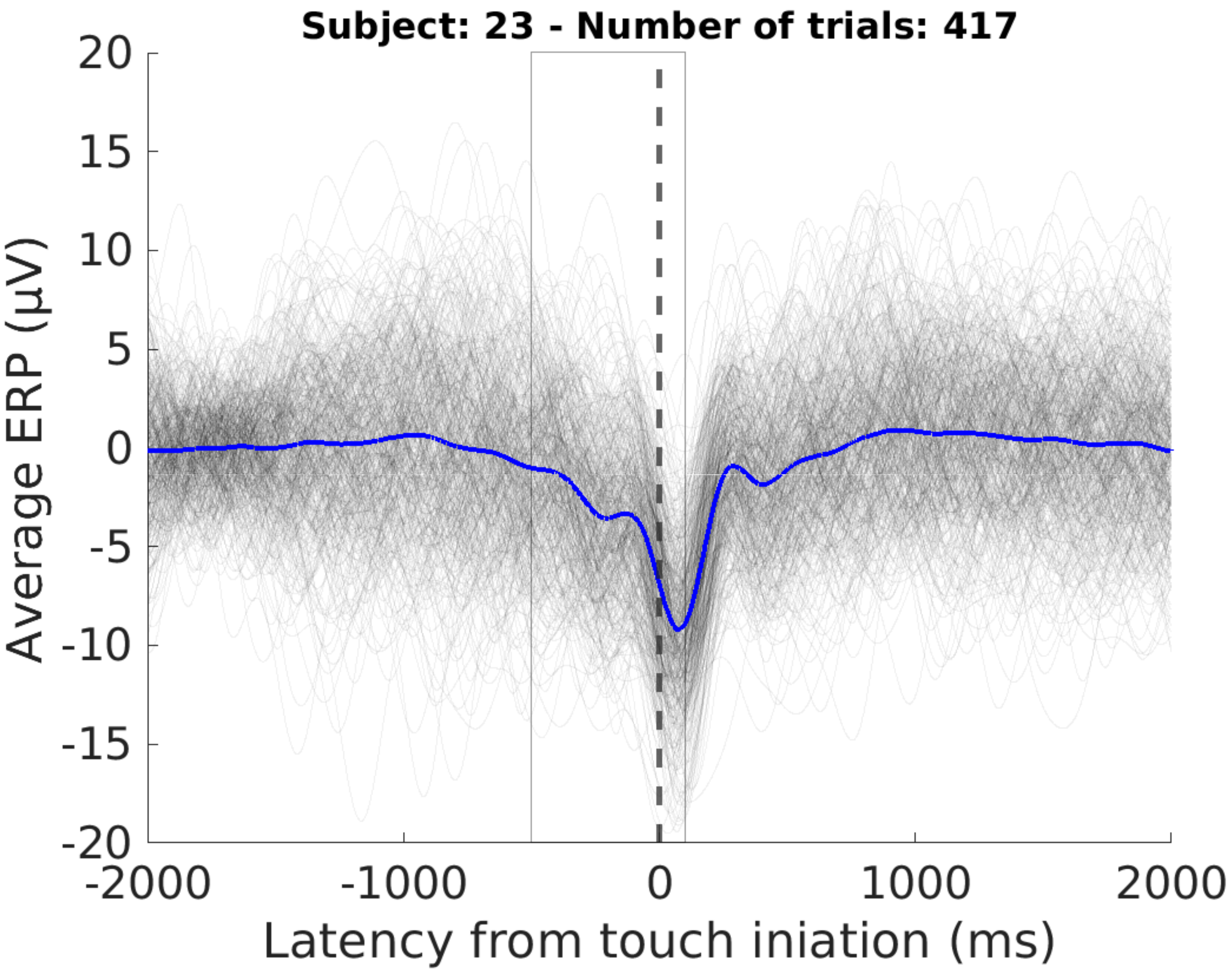

Subject: 23 - Number of trials: 417
Average ERP (µV)
20
15
10
5
0
-5
-10
-15
-20
-2000
-1000
0
1000
2000
Latency from touch iniation (ms)
