## Supplementary figures and images for "Multiple forms of neural processing when repeating voluntary thumb flexions"

### Supplementary Figure 2

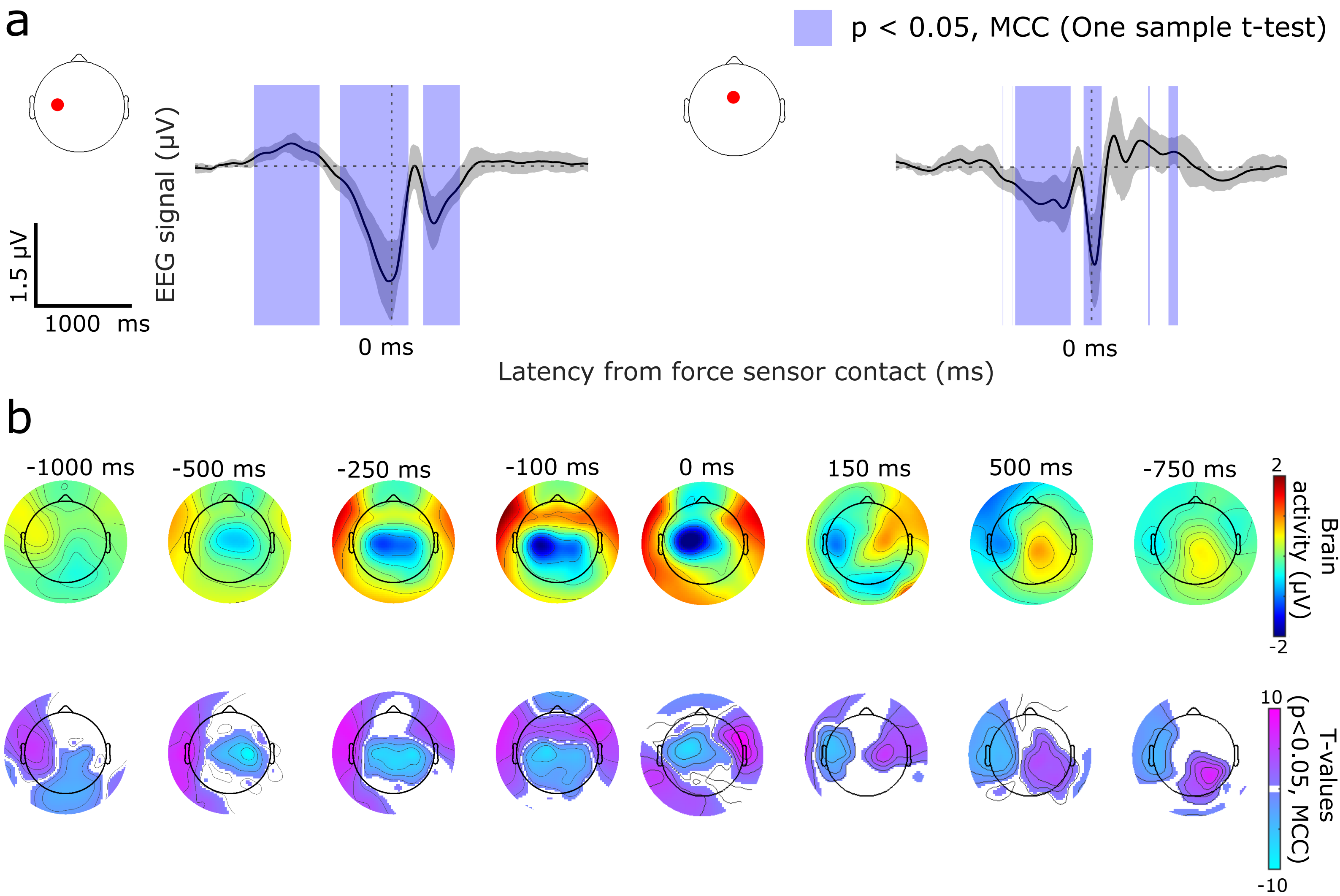

### Supplementary Figure 4

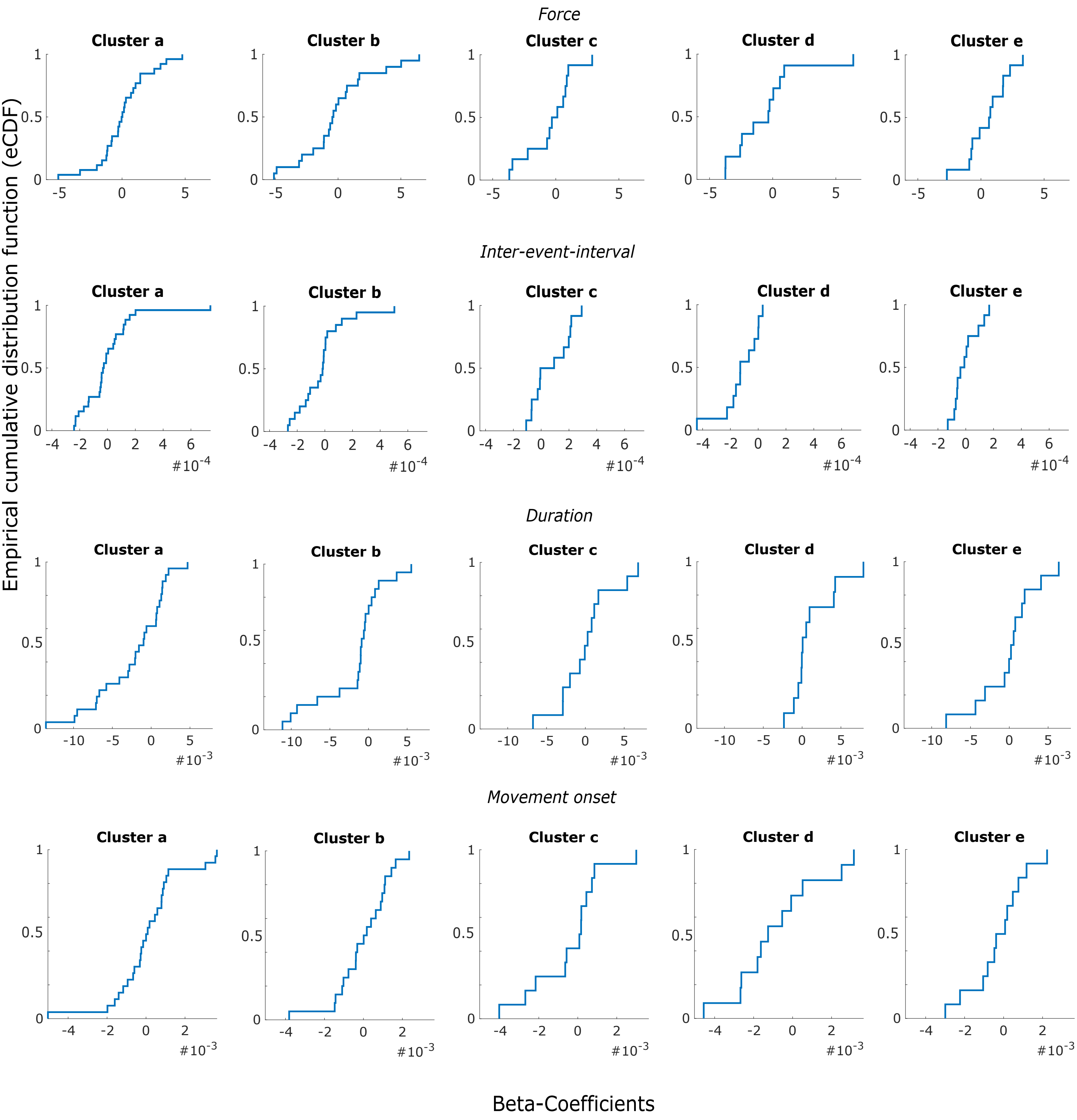

### Supplementary Figure 5

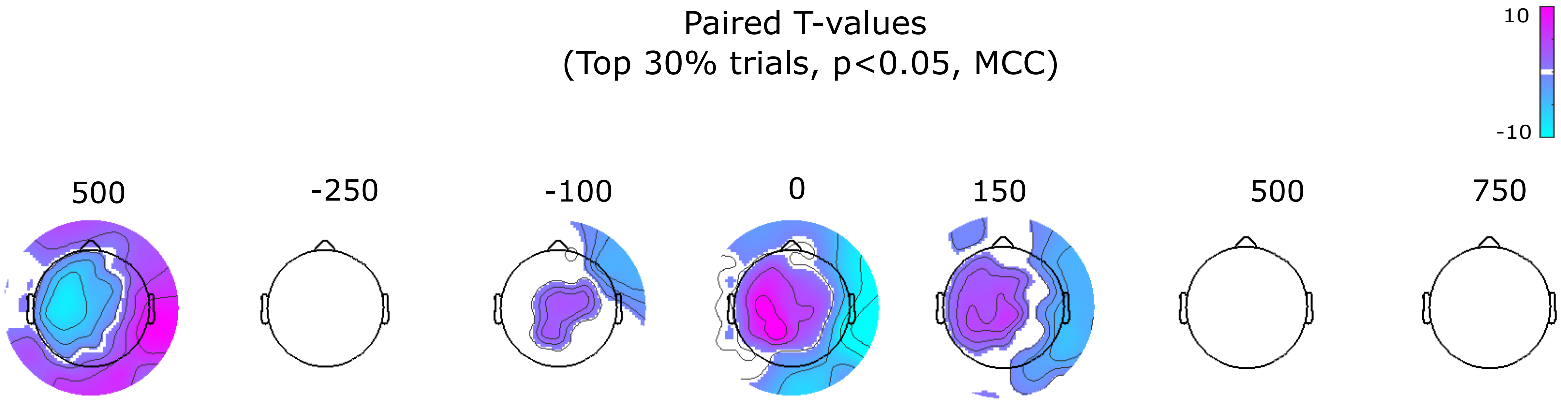

### Supplementary Figure 6

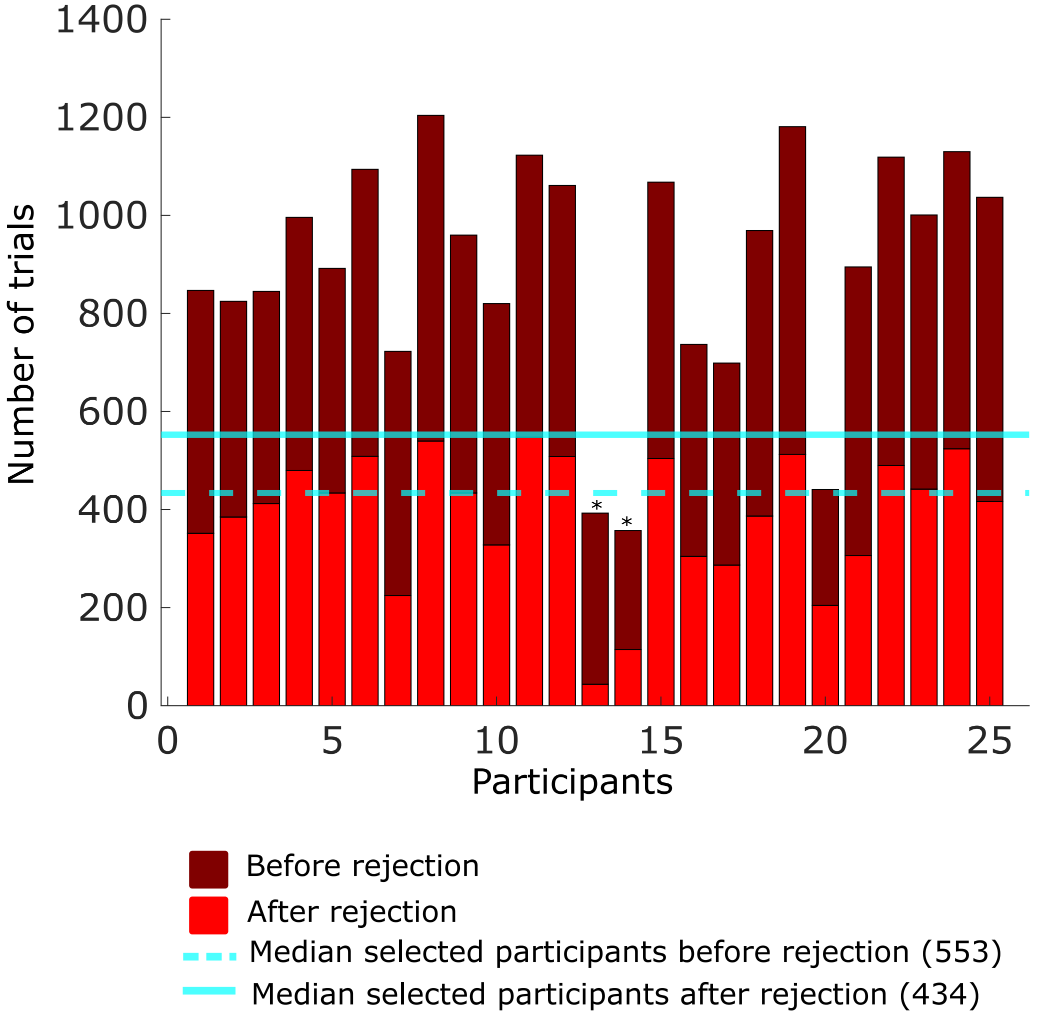


* Removed participants with less than 200 trials after rejection
