## Supplementary Figure 3 for "Multiple forms of neural processing when repeating voluntary thumb flexions"

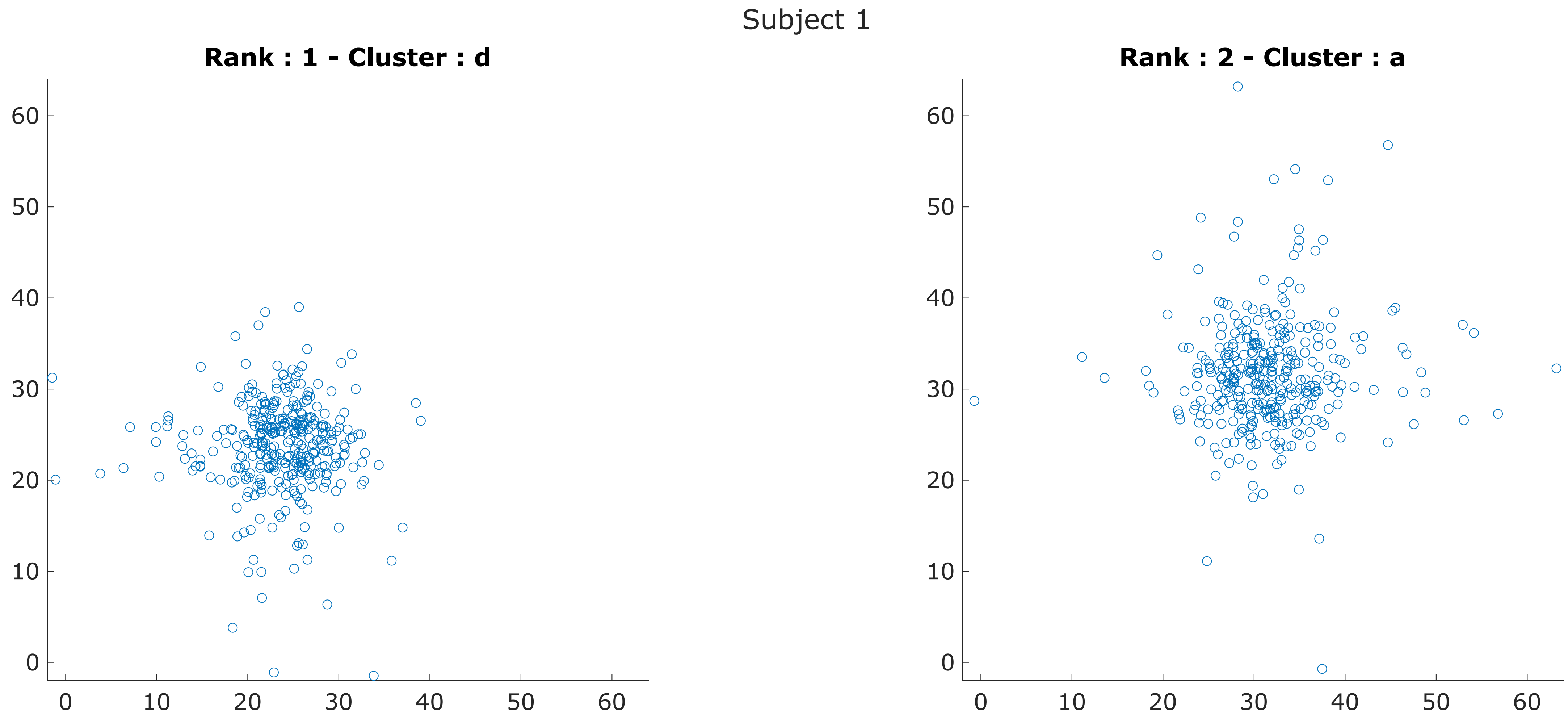
Subject 1
Rank : 1 - Cluster : d
Rank : 2 - Cluster : a
0
10
20
30
40
50
60

Subject 2

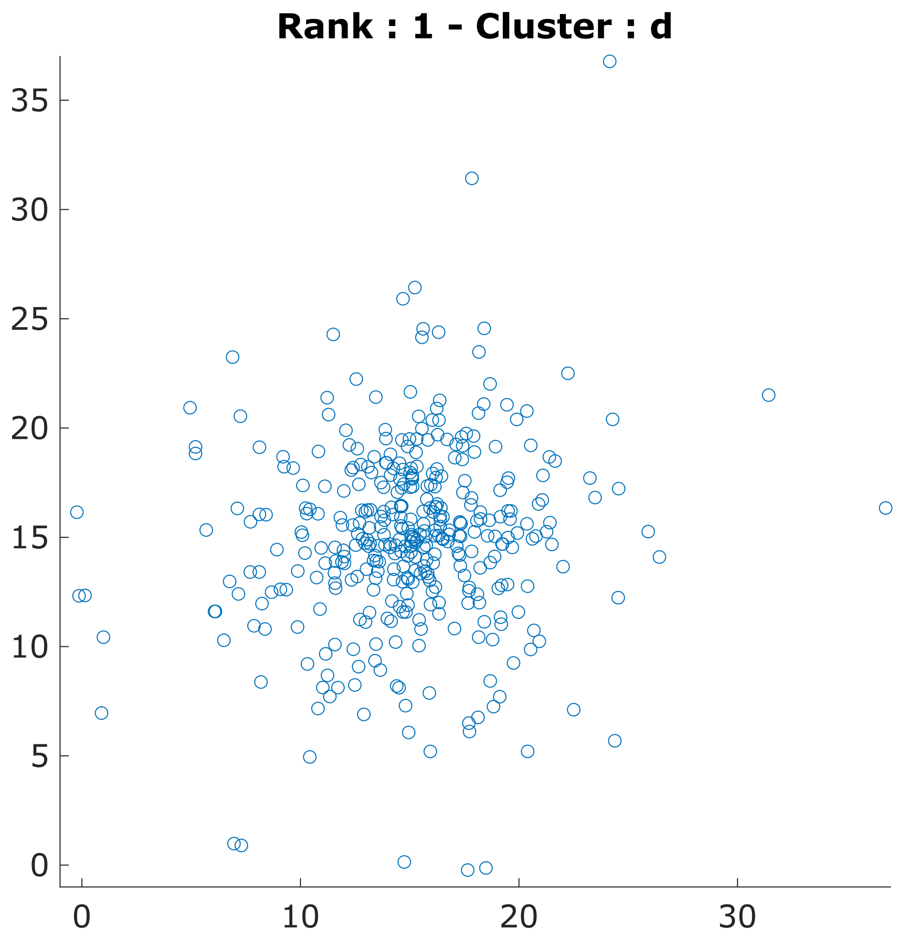
Rank : 1 - Cluster : d
35
30
25
20
15
10
5
0
0
10
20
30

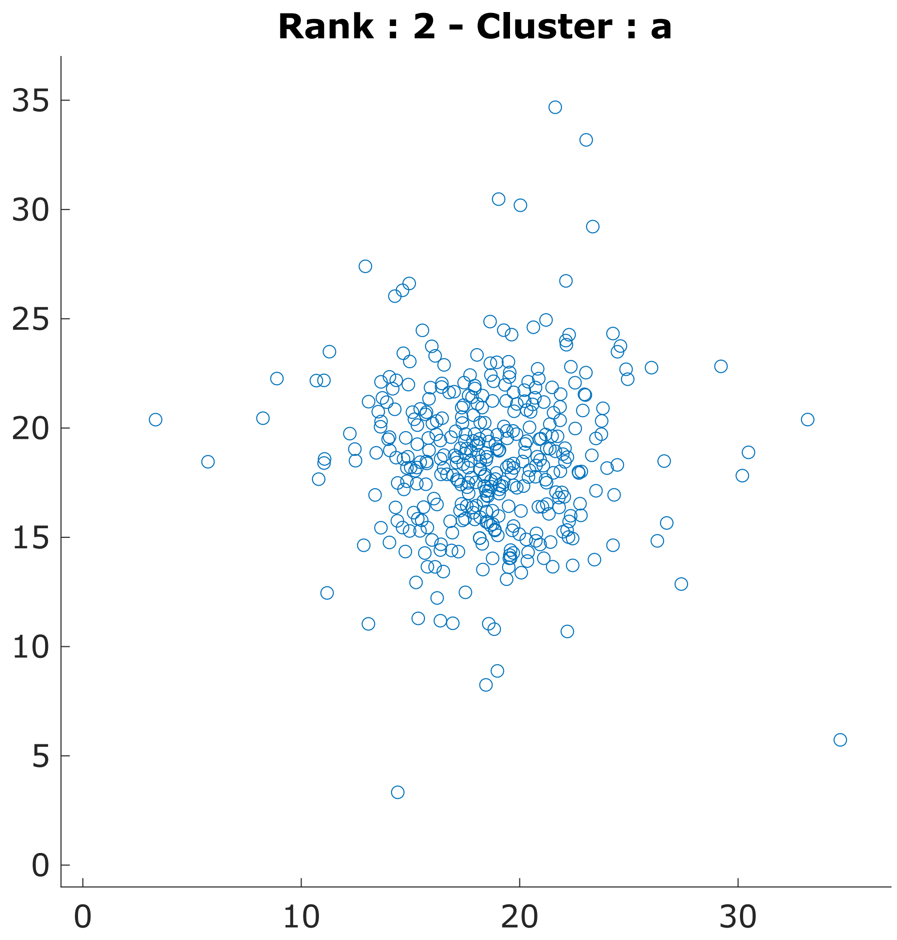
Rank : 2 - Cluster : a
35
30
25
20
15
10
5
0
0
10
20
30

Subject 3

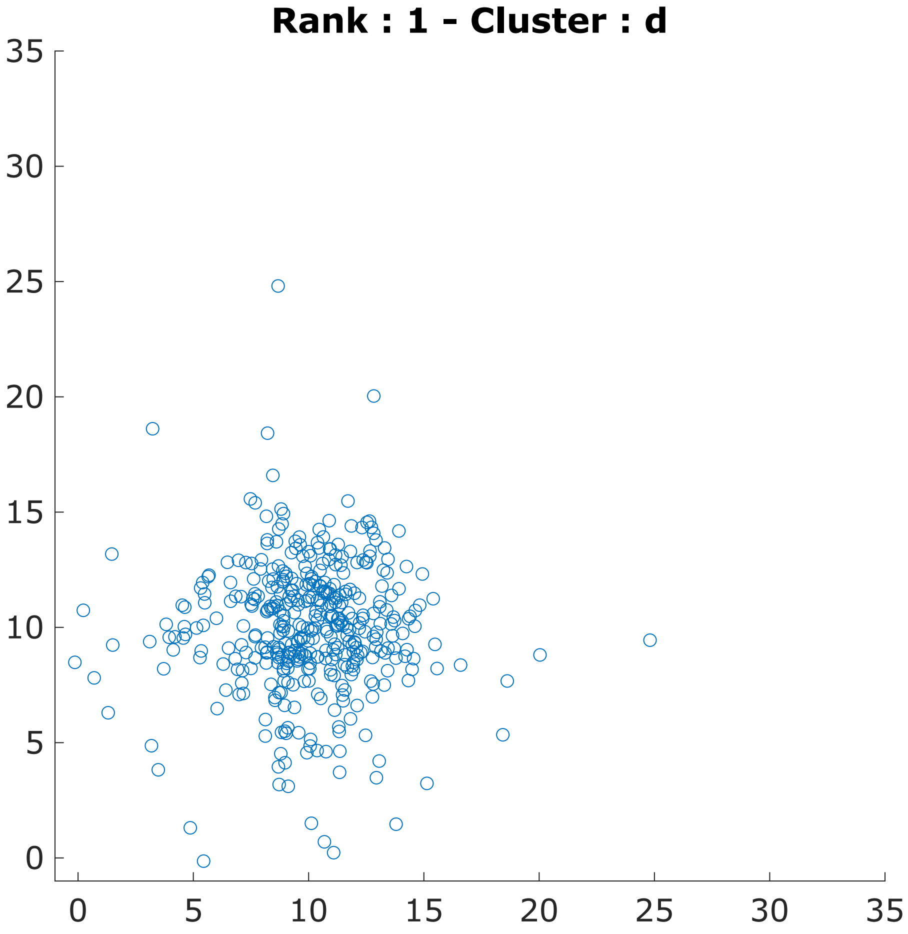
Rank : 1 - Cluster : d
0
5
10
15
20
25
30
35
0
5
10
15
20
25
30
35

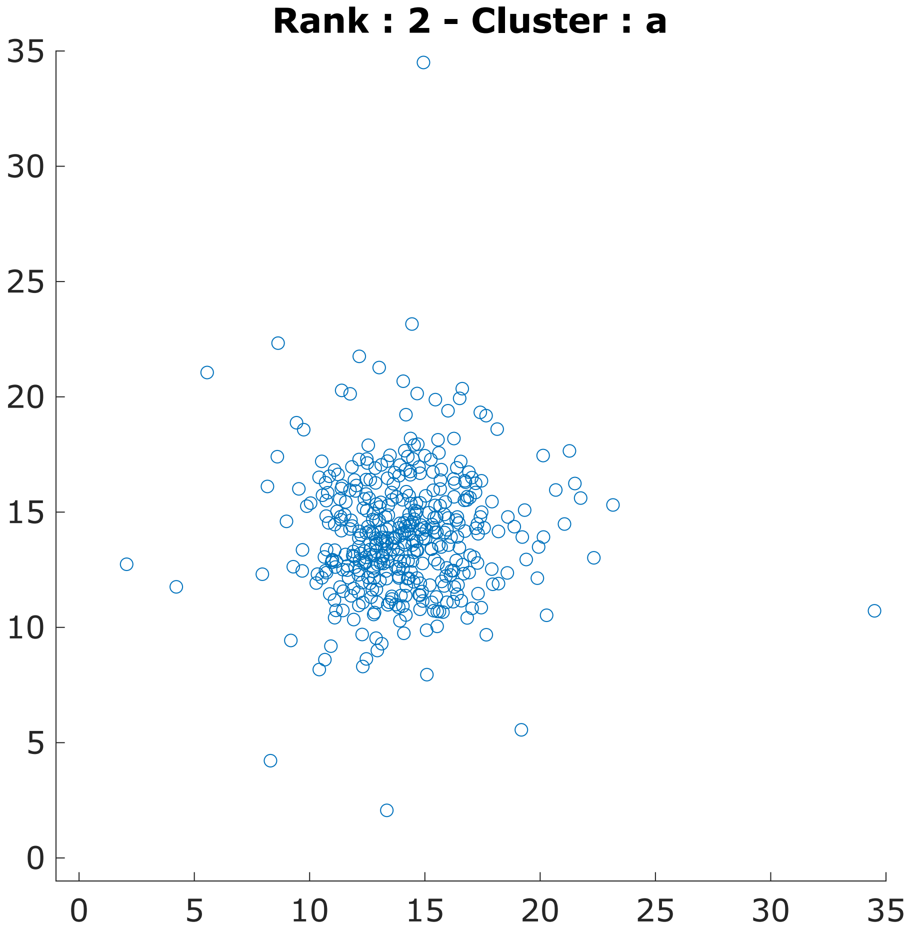
Rank : 2 - Cluster : a
0
5
10
15
20
25
30
35
0
5
10
15
20
25
30
35

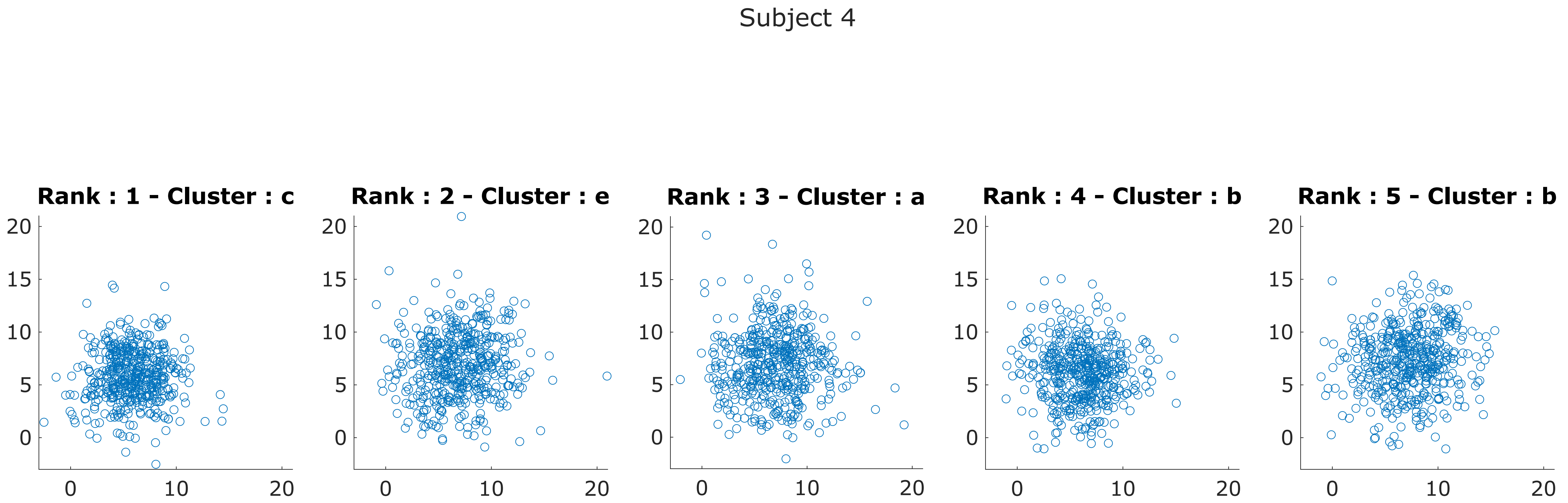

Subject 4
Rank : 1 - Cluster : c
Rank : 2 - Cluster : e
Rank : 3 - Cluster : a
Rank : 4 - Cluster : b
Rank : 5 - Cluster : b
0
5
10
15
20

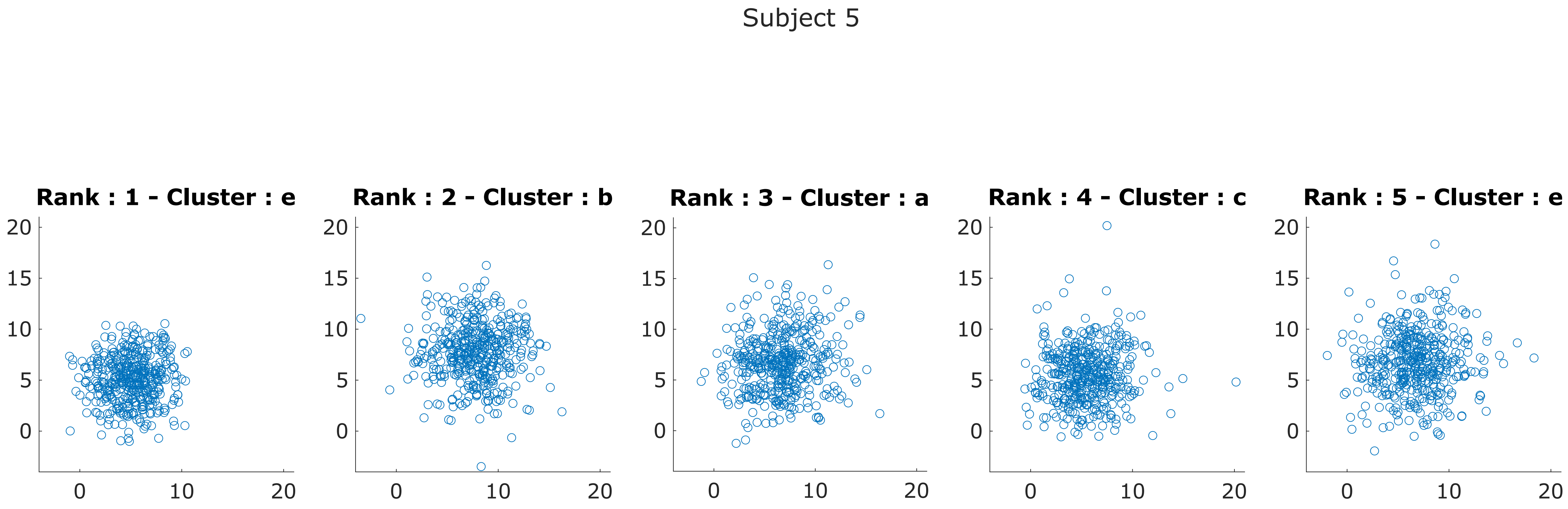
Subject 5
Rank : 1 - Cluster : e
Rank : 2 - Cluster : b
Rank : 3 - Cluster : a
Rank : 4 - Cluster : c
Rank : 5 - Cluster : e

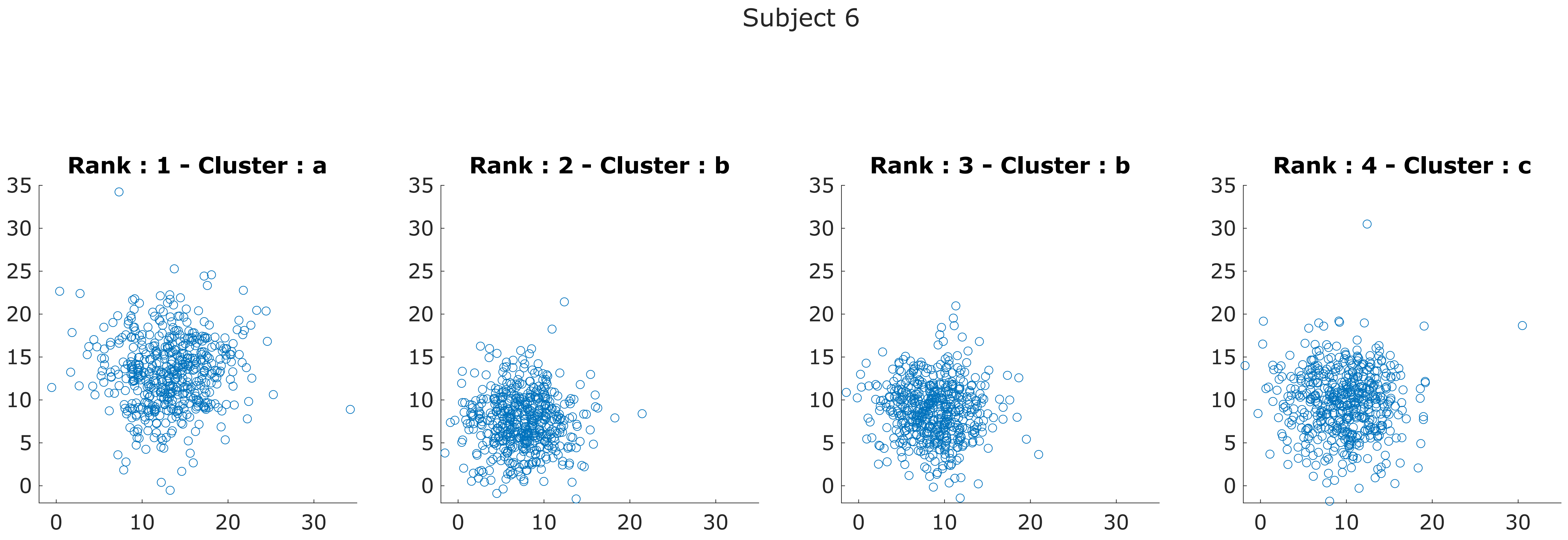
Subject 6
Rank : 1 - Cluster : a
Rank : 2 - Cluster : b
Rank : 3 - Cluster : b
Rank : 4 - Cluster : c
35
30
25
20
15
10
5
0
0
10
20
30

Subject 7

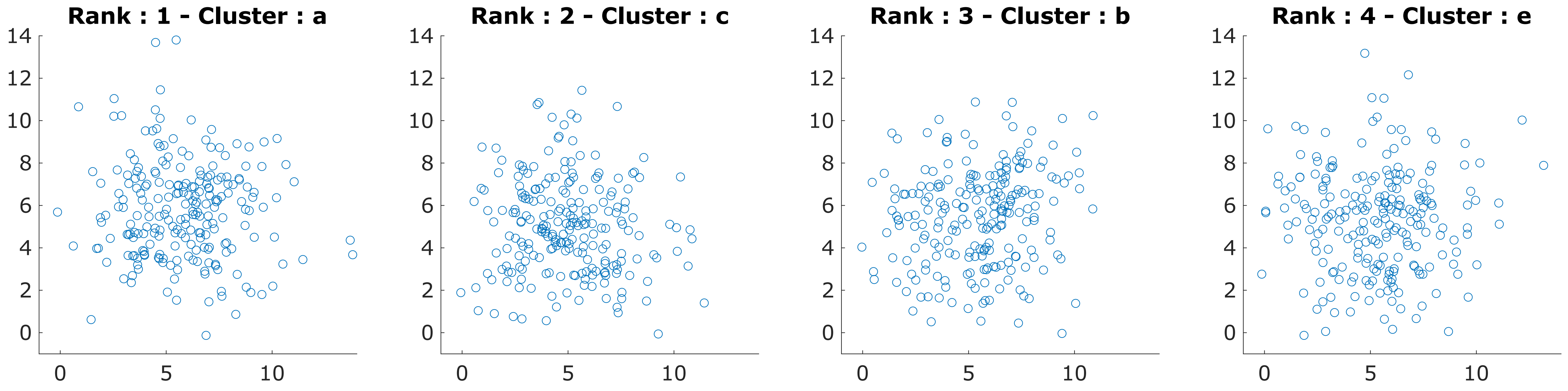
Rank : 1 - Cluster : a
Rank : 2 - Cluster : c
Rank : 3 - Cluster : b
Rank : 4 - Cluster : e
0
2
4
6
8
10
12
14
0
5
10

Subject 8

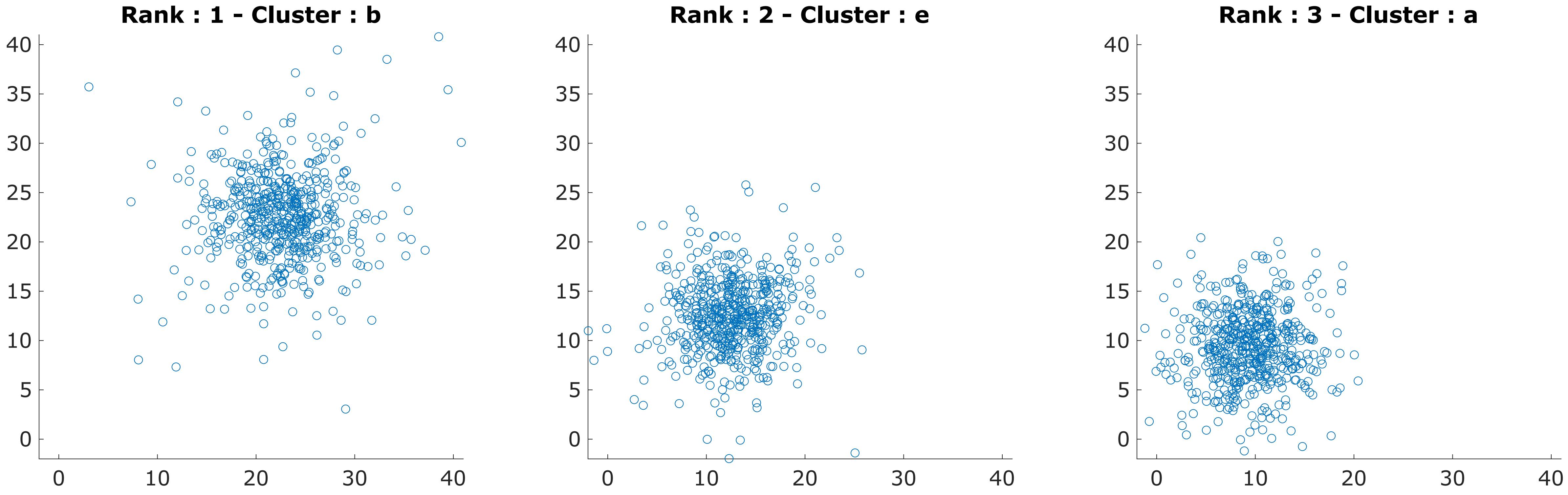
Rank : 1 - Cluster : b
Rank : 2 - Cluster : e
Rank : 3 - Cluster : a
0
5
10
15
20
25
30
35
40

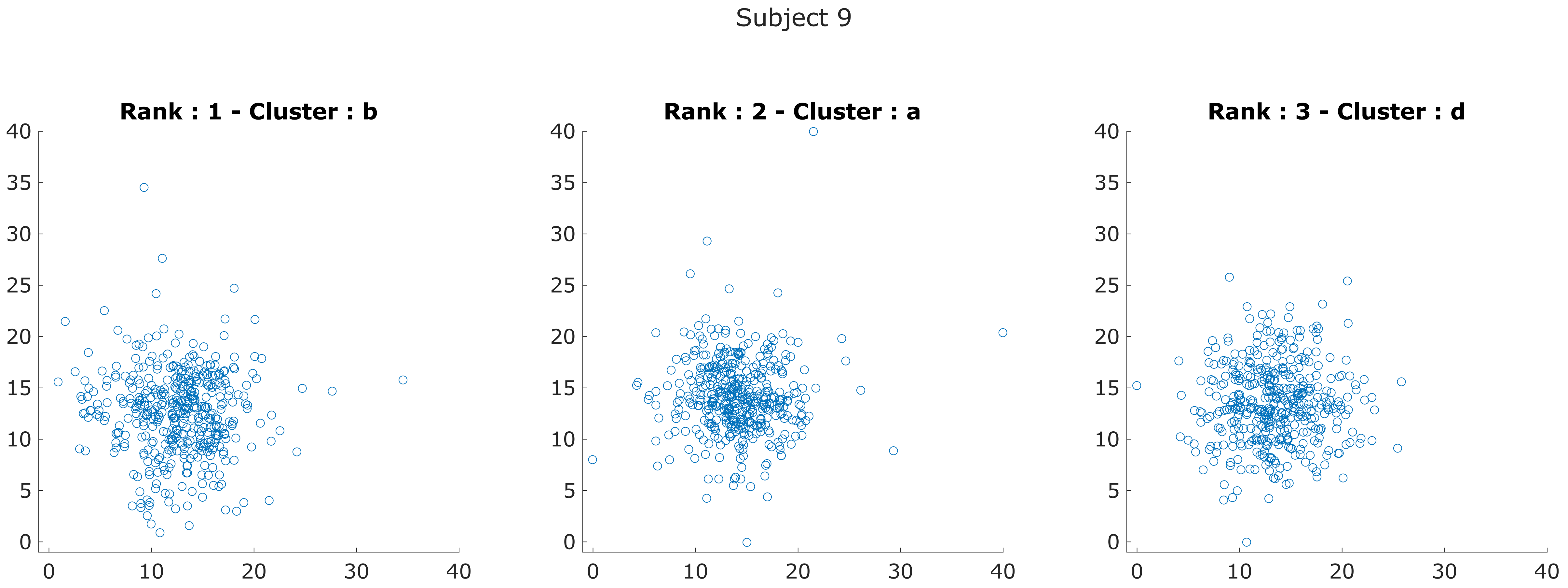
Subject 9
Rank : 1 - Cluster : b
Rank : 2 - Cluster : a
Rank : 3 - Cluster : d
0
5
10
15
20
25
30
35
40

Subject 10

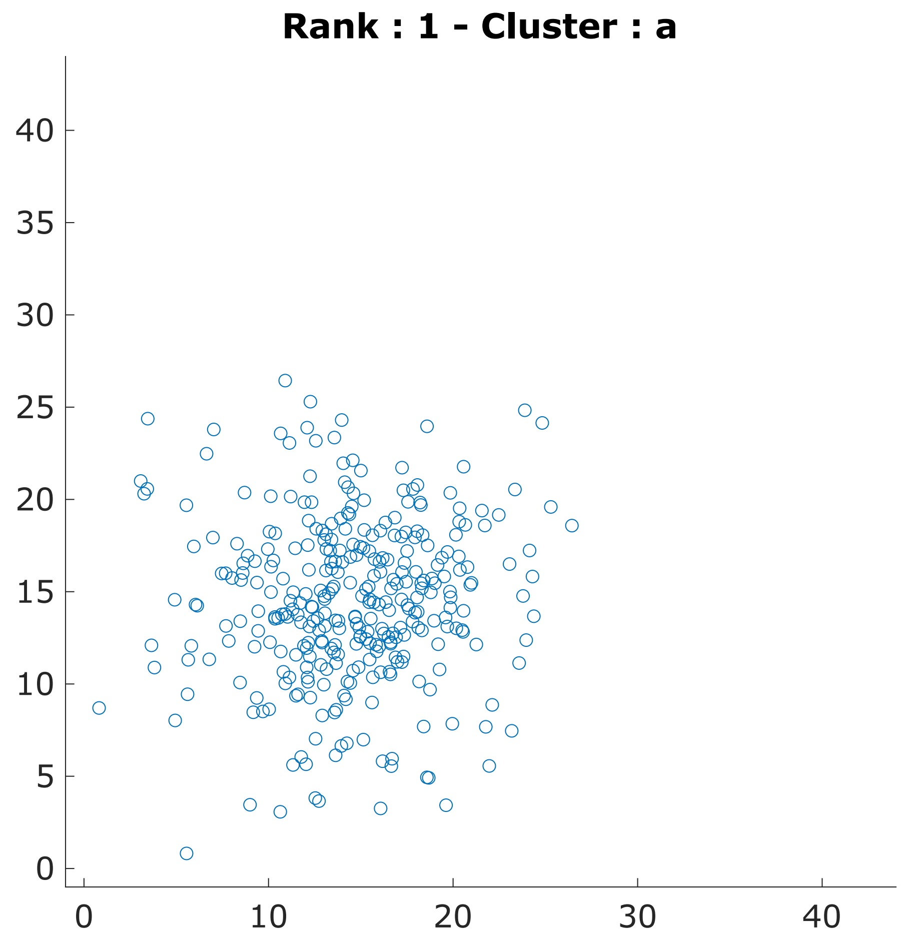

Rank : 1 - Cluster : a
0
5
10
15
20
25
30
35
40
0
10
20
30
40


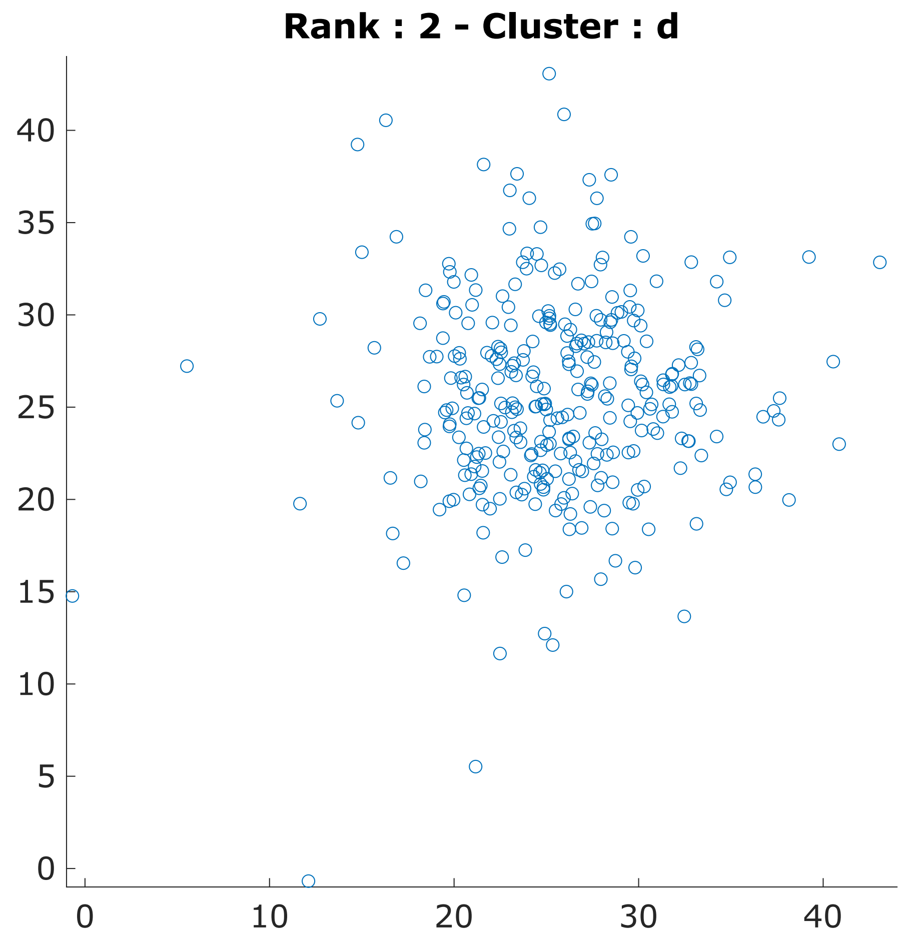

Rank : 2 - Cluster : d
0
5
10
15
20
25
30
35
40
0
10
20
30
40

Subject 11

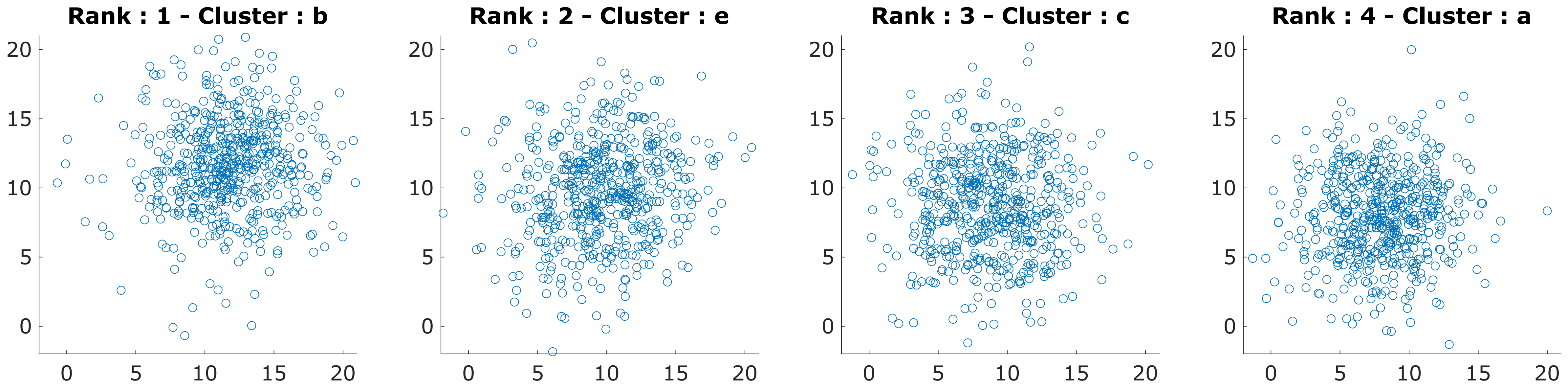
Rank : 1 - Cluster : b
Rank : 2 - Cluster : e
Rank : 3 - Cluster : c
Rank : 4 - Cluster : a

Subject 12

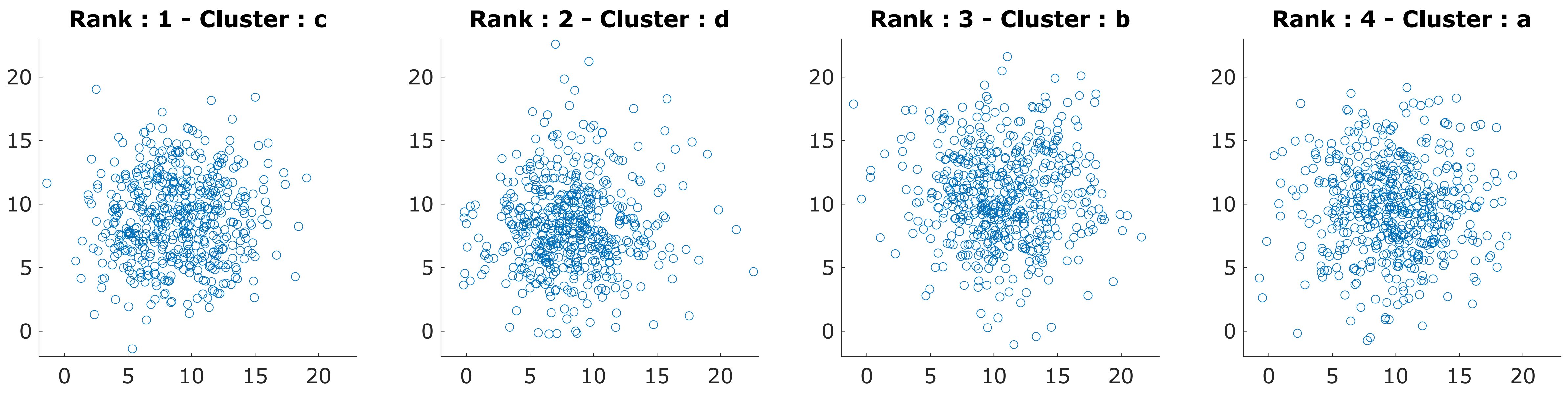
Rank : 1 - Cluster : c
Rank : 2 - Cluster : d
Rank : 3 - Cluster : b
Rank : 4 - Cluster : a
0
5
10
15
20

Subject 13

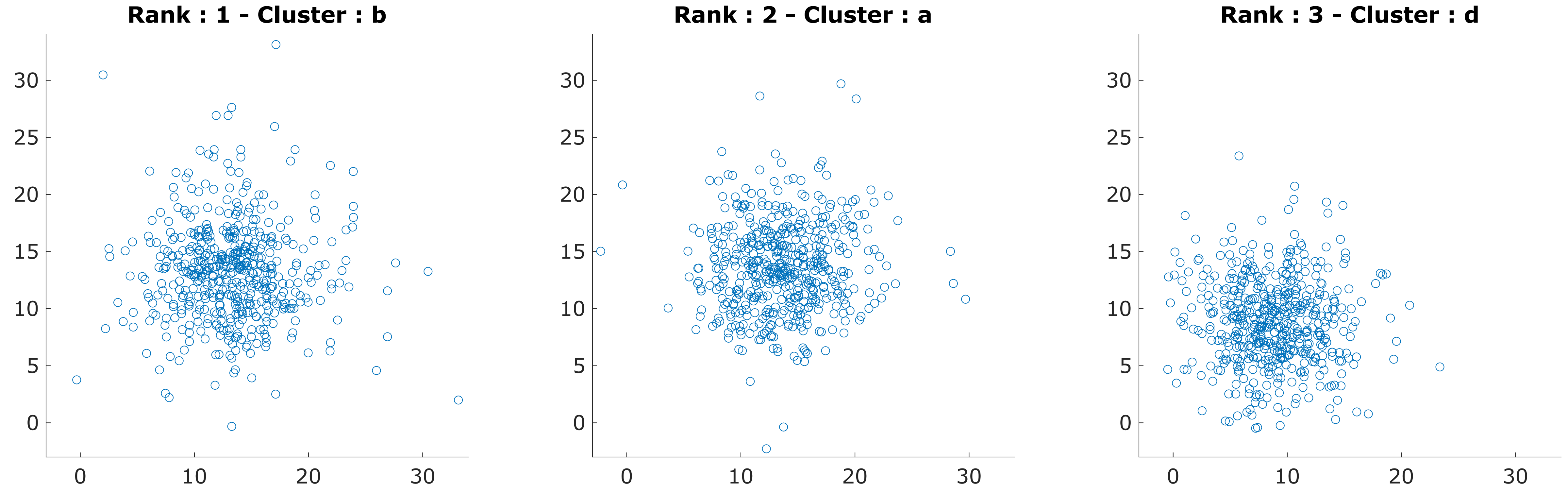

Rank : 1 - Cluster : b
Rank : 2 - Cluster : a
Rank : 3 - Cluster : d
0
5
10
15
20
25
30

### Subject 14

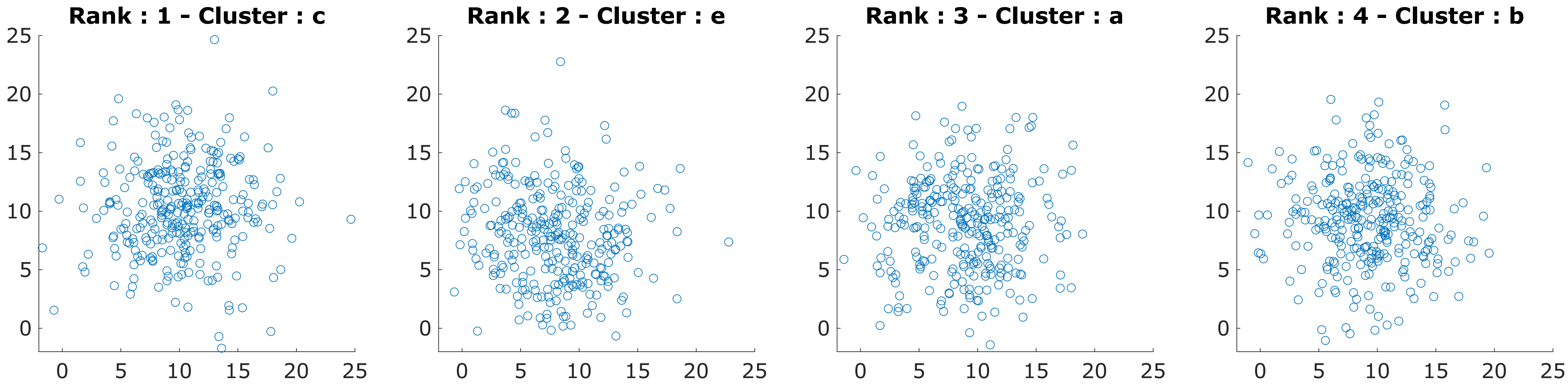

Rank : 1 - Cluster : c
Rank : 2 - Cluster : e
Rank : 3 - Cluster : a
Rank : 4 - Cluster : b
25
20
15
10
5
0
0 5 10 15 20 25

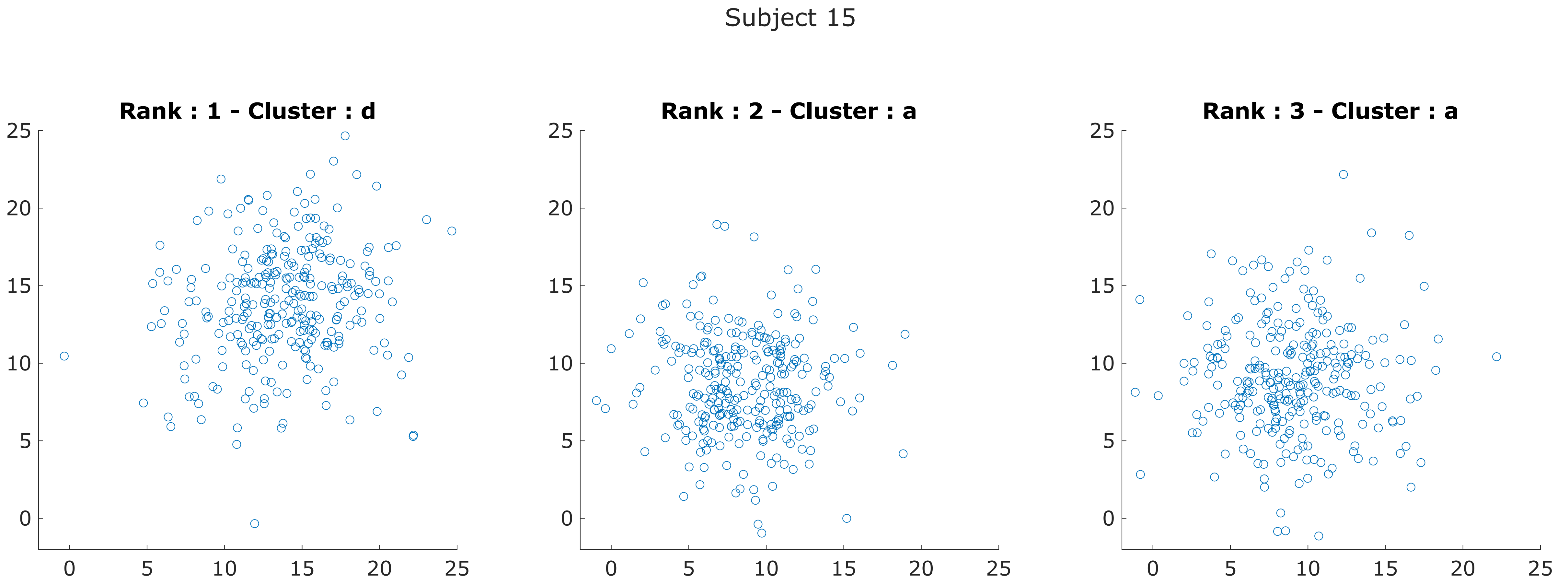
Subject 15
Rank : 1 - Cluster : d
Rank : 2 - Cluster : a
Rank : 3 - Cluster : a
25
20
15
10
5
0
0
5
10
15
20
25

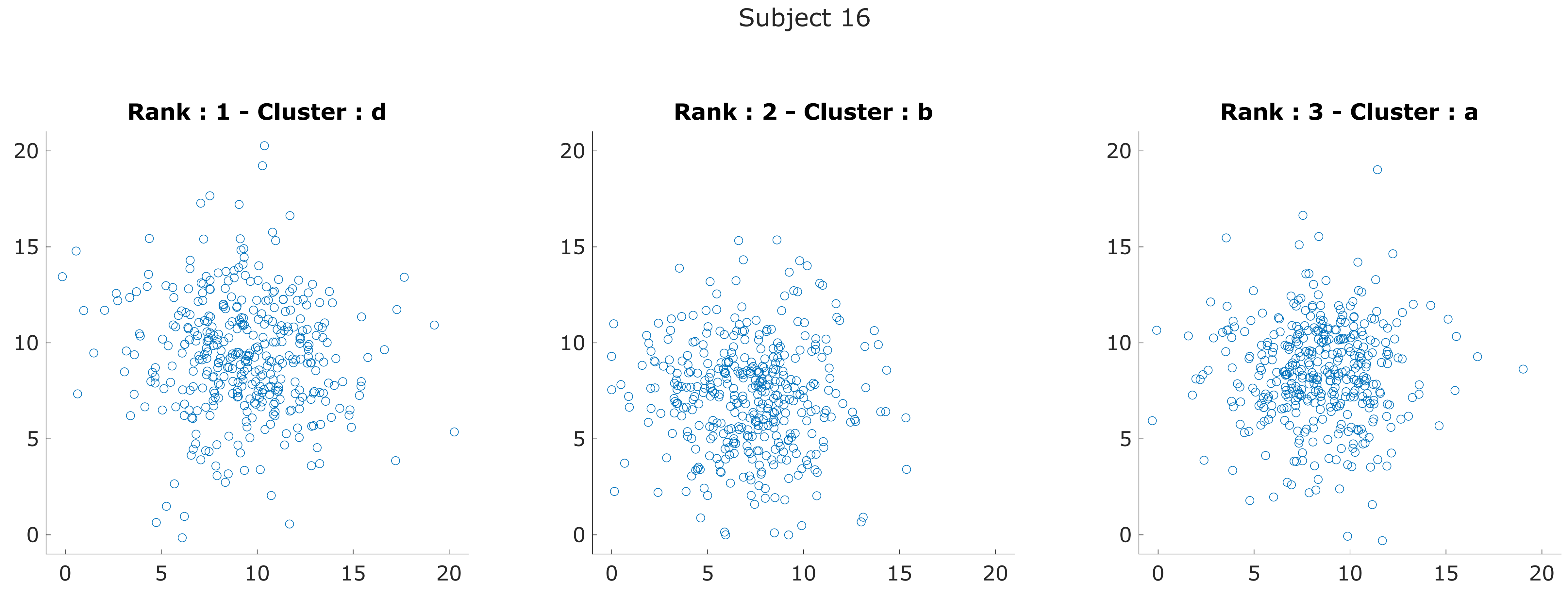
Subject 16
Rank : 1 - Cluster : d
Rank : 2 - Cluster : b
Rank : 3 - Cluster : a
0
5
10
15
20

### Subject 17

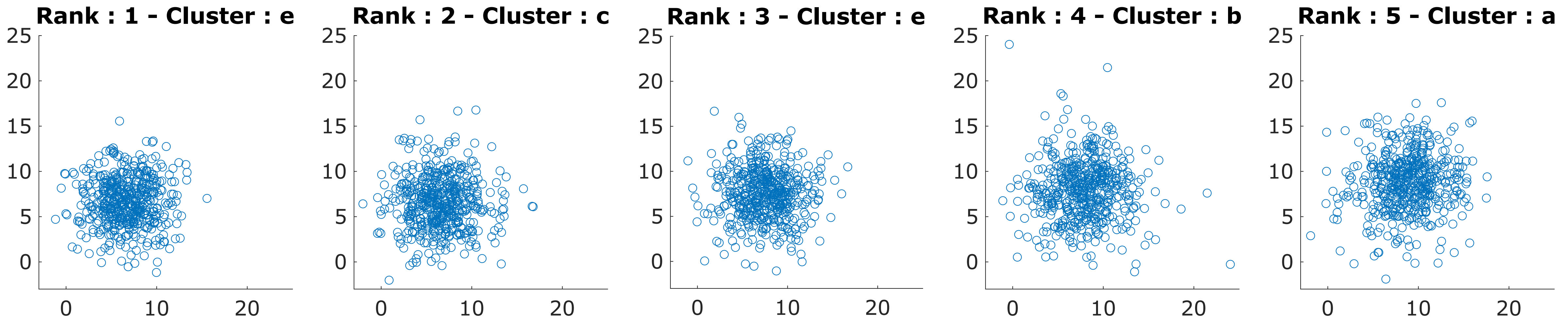
Rank : 1 - Cluster : e
Rank : 2 - Cluster : c
Rank : 3 - Cluster : e
Rank : 4 - Cluster : b
Rank : 5 - Cluster : a

Subject 18

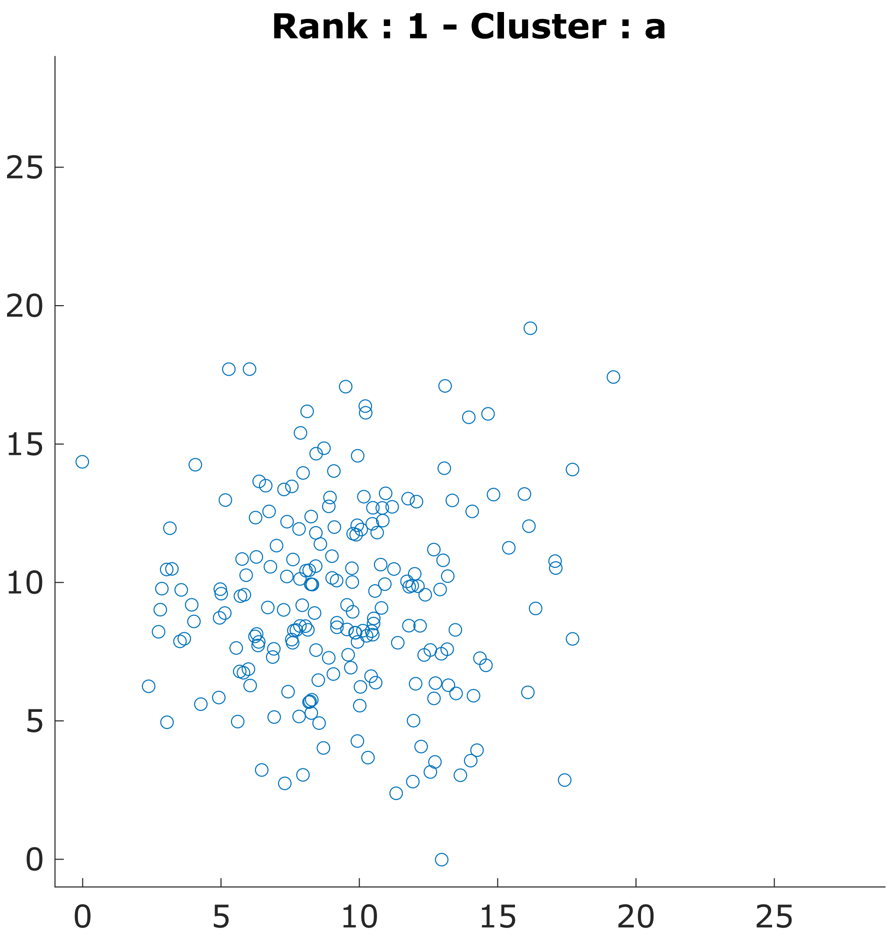


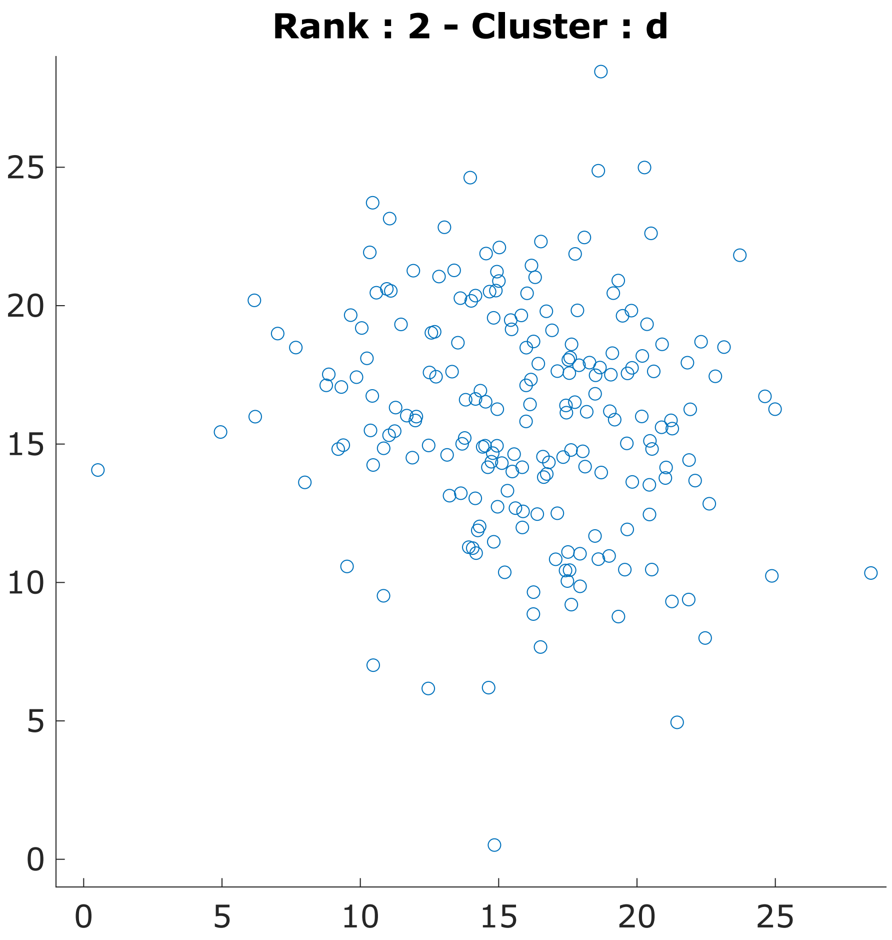

Subject 19

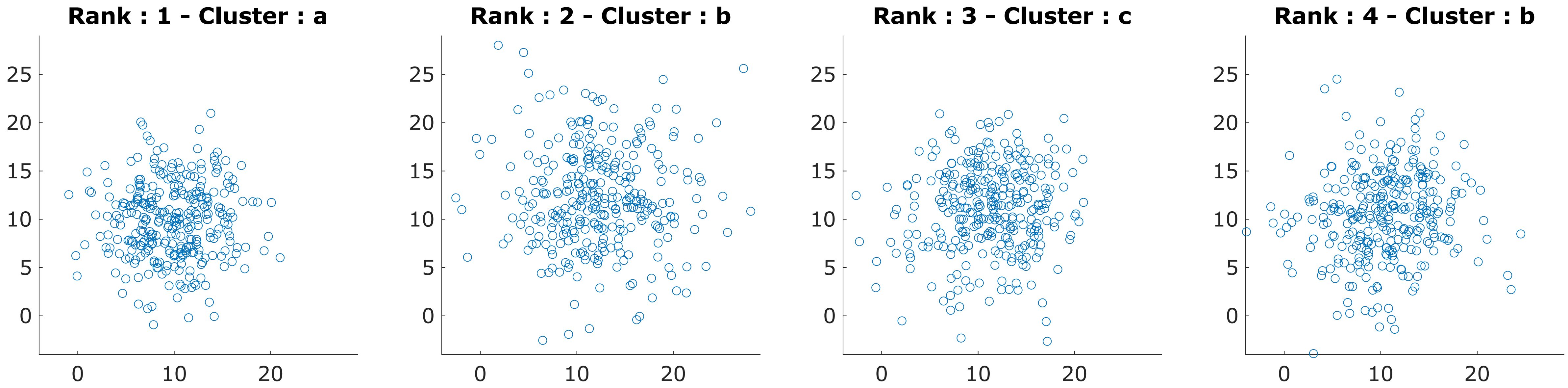
Rank : 1 - Cluster : a
Rank : 2 - Cluster : b
Rank : 3 - Cluster : c
Rank : 4 - Cluster : b
25
20
15
10
5
0
0
10
20

Subject 20

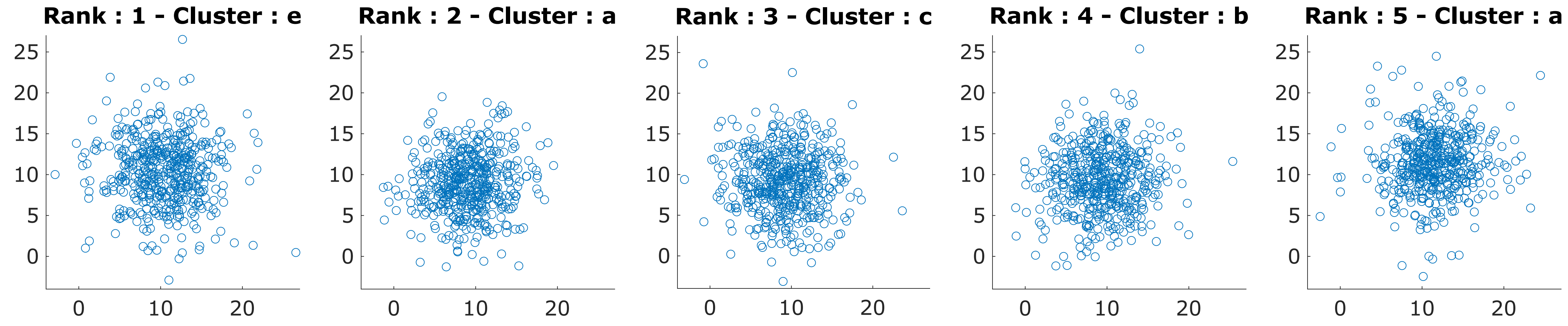

Rank : 1 - Cluster : e
Rank : 2 - Cluster : a
Rank : 3 - Cluster : c
Rank : 4 - Cluster : b
Rank : 5 - Cluster : a

Subject 21

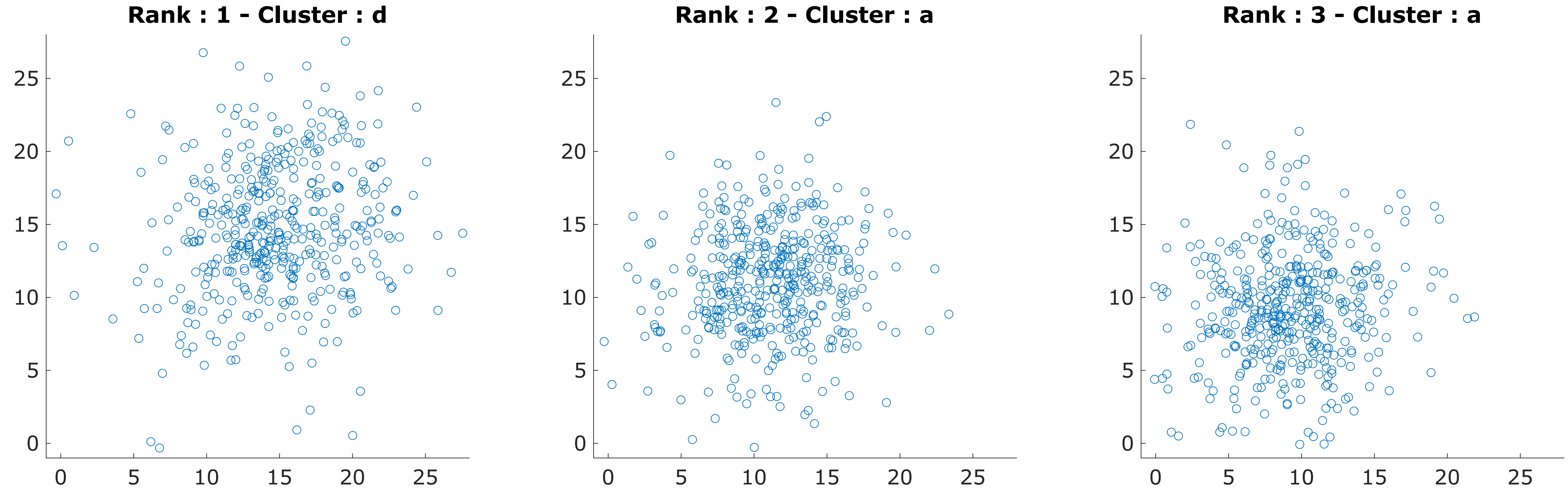

Subject 22

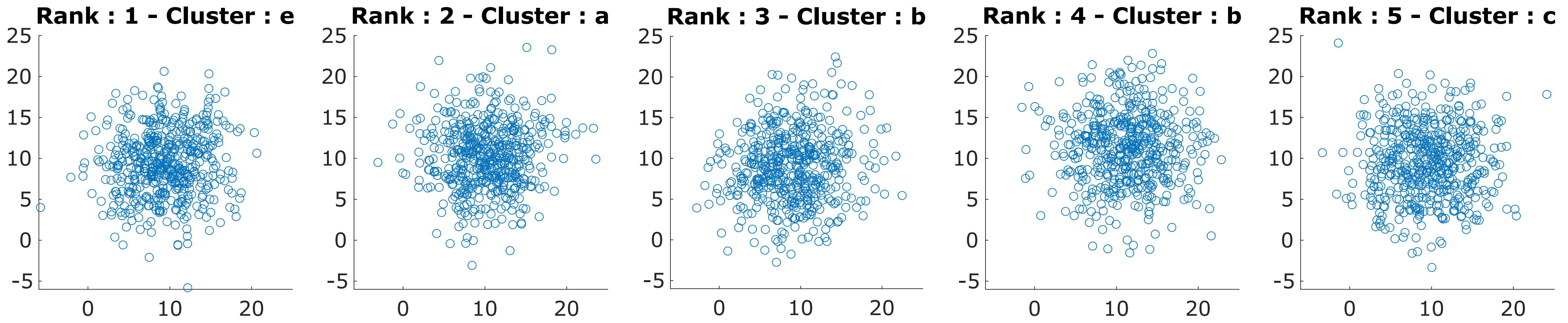
Rank : 1 - Cluster : e
Rank : 2 - Cluster : a
Rank : 3 - Cluster : b
Rank : 4 - Cluster : b
Rank : 5 - Cluster : c
25
20
15
10
5
0
-5
0
10
20

Subject 23

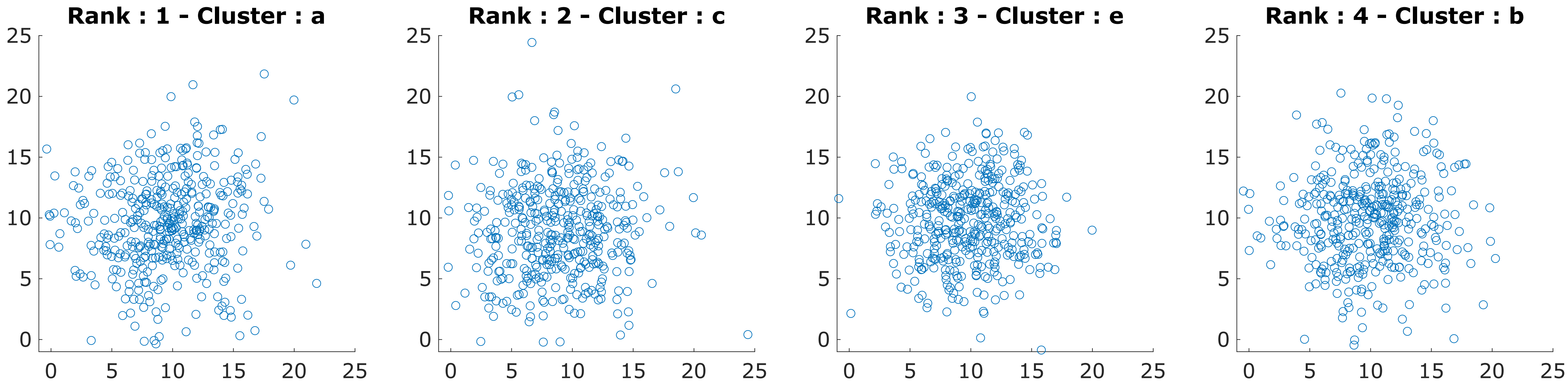

Rank : 1 - Cluster : a
Rank : 2 - Cluster : c
Rank : 3 - Cluster : e
Rank : 4 - Cluster : b
0
5
10
15
20
25
